# Feline coronavirus drug inhibits the main protease of SARS-CoV-2 and blocks virus replication

**DOI:** 10.1101/2020.05.03.073080

**Authors:** Wayne Vuong, Muhammed Bashir Khan, Conrad Fischer, Elena Arutyunova, Tess Lamer, Justin Shields, Holly A. Saffran, Ryan T. McKay, Marco J. van Belkum, Michael Joyce, Howard S. Young, D. Lorne Tyrrell, John C. Vederas, M. Joanne Lemieux

## Abstract

The COVID-19 pandemic, attributed to the SARS-CoV-2 coronavirus infection, resulted in millions infected worldwide and an immediate need for antiviral treatments. The main protease (M^pro^) in SARS-CoV-2 is a viable drug target because of its essential role in the cleavage of the virus polypeptide and subsequent viral replication. Feline infectious peritonitis, a fatal infection in cats caused by a coronavirus, was successfully treated previously with a dipeptide-based protease inhibitor. Here we show this drug, GC376, and its analog GC373, are effective inhibitors of the M^pro^ from both SARS-CoV and SARS-CoV-2 with IC_50_ values in the nanomolar range. Crystal structures of the SARS-CoV and SARS-CoV-2 M^pro^ with these inhibitors have a covalent modification of the nucleophilic Cys145. NMR analysis reveals that inhibition proceeds via reversible formation of a hemithioacetal. GC373 and GC376 are potent inhibitors of SARS-CoV-2 in cell culture, with EC_50_ values near one micromolar and little to no toxicity. These protease inhibitors are soluble, non-toxic, and bind reversibly. They are strong drug candidates for the treatment of human coronavirus infections because they have already been successful in animals (cats). The work here lays the framework for their use in human trials for the treatment of COVID-19.

---

The COVID-19 outbreak evolved into a pandemic due to the virulent nature of SARS-CoV-2, reaching over 3 million cases worldwide by end of April 2020, with the number of infected growing rapidly worldwide^1^. This current scenario contrasts the less virulent SARS outbreak in 2002-03, which had only 8000 cases and 774 deaths (WHO, 2004) ^2^. There is an urgent need for antiviral therapies for acute COVID-19 infections, especially until an efficacious vaccine is developed. The main protease of the coronavirus is a strong drug target due to its essential nature for virus maturation and subsequent infection^3^. Coronaviruses are RNA viruses that hijack the host’s translational machinery to generate viral proteins. The viral RNA encodes two overlapping polyproteins: pp1a and pp1ab, which are 450 kD and 750 kD, respectively. The polyproteins need to be cleaved in order to release individual functional proteins for viral replication and transcription. Viral encoded proteases include the main protease (M^pro^), also called 3CL^pro^, and a papain-like protease (PL^pro^). M^pro^ cleaves the polyproteins at 11 positions primarily at conserved Leu Gln | Ser Ala Gly sequences, which allows for virus assembly. Given its crucial role in virus replication, the SARS-CoV-2 M^pro^ is a prominent drug target for COVID-19 antiviral therapy. The coronavirus M^pro^ is a cysteine protease for which many different inhibitor classes exist^4^. Protease inhibitors are common drug candidates if they meet the requirements of low toxicity, solubility, and reversibility^5^. Several proteases have been identified as molecular targets and used for the development of novel classes of drugs^5^ including Tipranavir for the treatment of HIV^6^. However, inhibition of cysteine proteases by thiol reactive species is often untenable for human drugs unless the inhibitor is reversible. Michael acceptor drugs that are irreversible *in vivo*, such as Rupintrivir, have failed in clinical trials due to low bioavailabilty^3^. Undesired irreversible reaction occurs with numerous mammalian thiols to destroy the inhibitor. Reaction with host protein thiols could also potentially lead to acute toxicity or immune reaction. In this regard, the reversible reaction of thiols with aldehyde inhibitors to make hemithioacetals presents a unique opportunity for effective cysteine protease inhibition, as they can potentially bind more effectively in the active site of their target protein than with other thiols^7^. Water soluble aldehyde bisulphite adducts are readily made, reversibly from the parent aldehyde under physiological conditions, and can be ideal prodrugs for cysteine protease inhibition as described below.

In early studies we developed peptide-based inhibitors, including aldehydes, against viral cysteine proteases^7^ that were subsequently studied with M^pro^ during the SARS coronavirus (SARS-CoV) outbreak in 2003^8^. Peptide aldehydes and their bisulphite derivatives were later used to inhibit the main protease of the Feline Coronavirus FCoV^9^. FCoV generally causes mild symptoms, but it can lead to feline infectious peritonitis (FIP), which is usually fatal in cats. The bisulphite adduct GC376, which converts readily to peptide aldehyde GC373, was well tolerated and able to reverse the infection in cats^10^. This, along with other studies that included ferret and mink coronavirus M^pro^, demonstrated the broad specificity of this protease inhibitor^11^. A crystal structure of GC376 was solved with the homologous MERS M^pro^ and demonstrated a covalent interaction with the catalytic cysteine of the M^pro^ ^12^. Recently, structures of the SARS-CoV-2 M^pro^ protease were solved with a peptide-based ketoamide inhibitor^13^ and various re-purposed drugs such as anti-cancer agents^14^. However, these M^pro^ inhibitors have not been tested in animal models of coronavirus infection nor have they been reported in human or animal trials for SARS. A number are known to have severe side effects and human cell toxicity, especially those that are anticancer agents. In this study, we examine whether GC373 and GC376, established as effective drugs in cats, inhibit SARS-CoV-2 M^pro^ reversibly and have potential for use as antiviral therapy in humans.

## GC373 and GC376 inhibit SARS-CoV-2 M^pro^

We synthesized the key dipeptidyl compounds, aldehyde GC373 and bisulphite adduct GC376 (**Figure 1A**)^9^ (**Extended data, Figure S1**), to test whether these FIP inhibitors are efficacious towards the M^pro^ of the SARS-CoV-2 and M^pro^ from SARS-CoV (associated with the 2002 outbreak). This compound consists of a glutamine surrogate in the substrate, P1 position, a Leu in P2 position and benzene ring in the P3 position, which reflects the known specificity for the SARS-CoV-2 M^pro^. The SARS-CoV-2 M^pro^ was cloned as a SUMO-tag fusion, which allowed for high-yield expression, enhanced stability, and generation of native N- and C-termini (**Extended data, Figure S2** and **S3)**. Similarly, SARS-CoV M^pro^ was expressed and purified to obtain native N- and C-termini according to previous methods^8^. Kinetic parameters for both the SARS-CoV M^pro^ and SARS-CoV-2-M^pro^ were determined using a synthetic peptide FRET-substrate with an anthranilate-nitrotyrosine donor-acceptor pair (Abz-SVTLQSG-Tyr^NO2^R - **Extended data, Figure S4**) as it displays over 10-fold more sensitivity compared to the equivalent EDANS-Dabcyl system^15^. Both SARS-CoV-2 M^pro^ and SARS-CoV M^pro^ exhibited cooperative substrate binding of the FRET-substrate (**Extended data, Figure S5**)

IC_50_ measurements revealed that both GC373 and GC376 inhibit the SARS-CoV M^pro^ and the SARS-CoV-2 M^pro^ *in vitro* at nanomolar concentrations (**Figure 1B and C**). For the SARS-CoV-2 M^pro^, IC_50_ for GC373 and GC376 are 0.40±0.05 μM and 0.19±0.04 μM, respectively. This is in agreement with studies of these compounds with M^pro^ from related viruses. For FCoV M^pro^ the IC_50_ for GC376 was 0.04 ±0.04 μM and for GC373 was 0.02 ±0.01 μM^9^. For SARS-CoV M^pro^ we observed an enhanced IC_50_, demonstrating the broad inhibition by both compounds, with GC373 and GC376 being 0.070 ± 0.02 μM and 0.05 ± 0.01 μM, respectively. The bisulphite adduct GC376 shows slightly higher potency for both enzymes compared to the free aldehyde. Our *in vitro* IC_50_ values for GC373 and GC376 reflect tight binding for the SARS-CoV-2 M^pro^ compared to other inhibitors tested *in vitro*, e.g. ebselen (IC_50_ 0.67 µM)^14^, tideglusib (IC_50_ 1.55 µM)^14^, carmofur (IC_50_ 1.82 µM)^14^, disulfiram (IC_50_ 9.35 µM)^14^, shikonin (IC_50_ 15.75 µM)^14^, PX-12 (IC_50_ 21.39 µM)^14^. Both GC373 and GC376 are also more potent than recently reported ketoamide inhibitors (IC_50_ of the lead compound 0.67 µM)^13^. Recently, a related peptidyl inhibitor was reported with a similar warhead to our compound, but with an indole group at the P3 position and an IC_50_ of 0.05±0.005 µM^16^. However, that compound has not been demonstrated to be efficacious in animals, as is the case for GC376.

**Figure 1.**
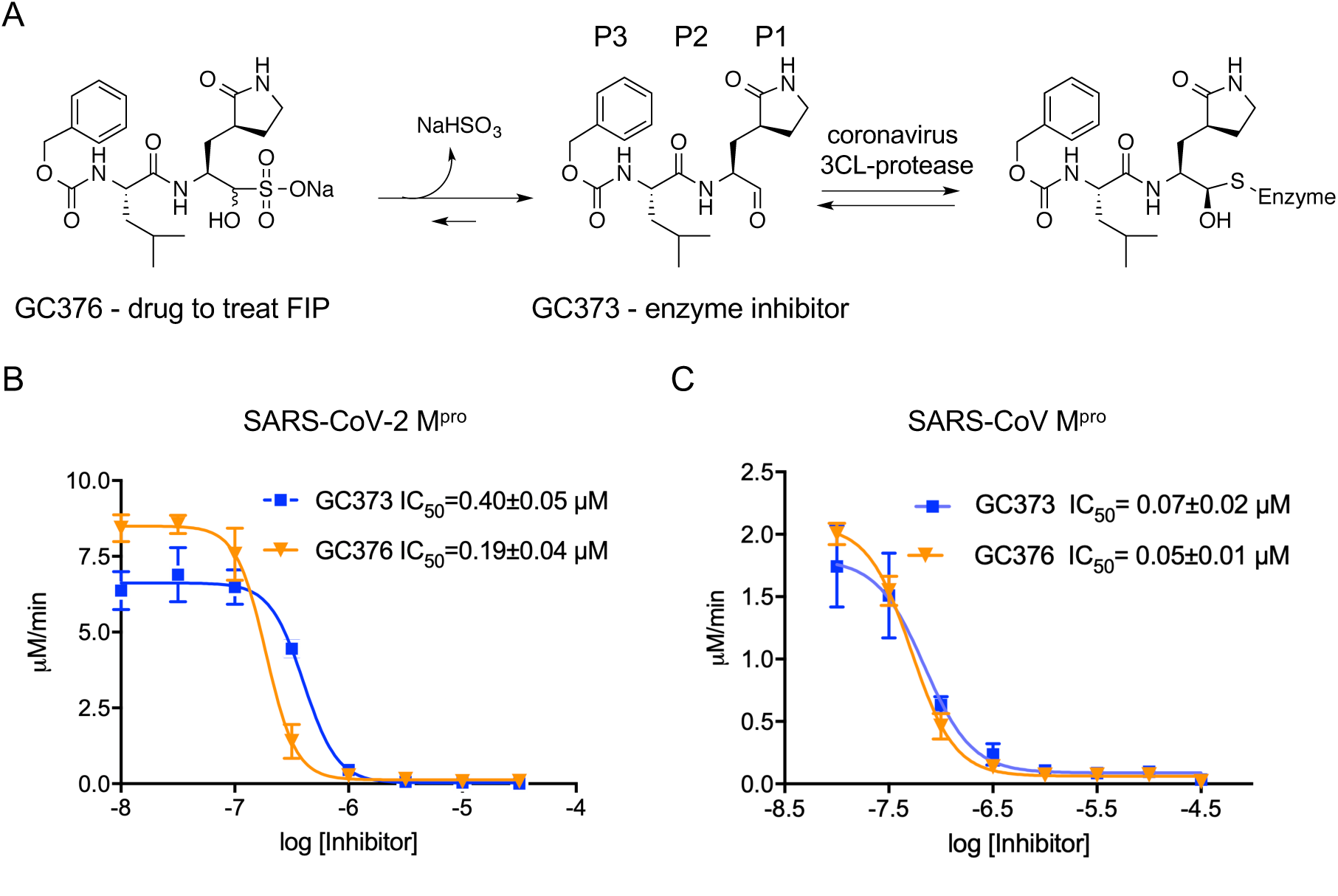
**A)** Schematic representation of inhibitor prodrug GC376, used to cure cats of FCoV, and GC373, the actual protease inhibitor. **B)** IC_50_ values for GC373 and GC376 for SARS-CoV-2-M^pro^ and **C)** SARS-CoV M^pro^ cleavage of Abz-SVTLQSG-Y(NO2)-R. N=3, values are represented as mean±SE.

## Crystal structure of SARS-CoV-2 M^pro^ in complex with GC373 and GC376

To gain insight into the mechanism of inhibition, the SARS-CoV-2 M^pro^ crystal structures with inhibitors GC373 and GC376 were determined at 2.0 Angstroms (**Figure 2 and Extended data, Table S1**). The three-dimensional structure of the SARS-CoV-2 M^pro^ (PDB Code 6WTM) is highly similar to the recently solved structures with an RMSD of 0.38 Å^2^ (PDB Code 6LU7) ^13, 14^. SARS-CoV-2 M^pro^ crystallized as a dimer facilitated by an N-finger of protomer A (residues 1 to 7) that fits into a pocket in protomer B. Each promoter displayed a two-lobe structure with one lobe composed of two-antiparallel β-barrels (Domains I and II), which form a chymotrypsin and 3C-like peptidase fold, with the active site comprised of a Cys-144 and His-41 dyad located at the domain interface. The oxyanion hole, influenced by dimerization^17^, is formed from the main chain residues Gly143, Ser144, and Cys145. The C-terminal domain III, is involved in domain swapping and facilitates dimer formation^18^. Molecular replacement with structure 6Y7M.PDB revealed electron density in the Fo-Fc map at the catalytic cysteine for both inhibitors GC373 (PDB Code 6WTK) and GC376 (PDB Code 6WTJ). In both structures the peptidyl inhibitor is covalently attached to Cys-145 as a hemithioacetal, showing that as expected the bisulphite group leaves GC376 (**Extended data, Figure S6**). In contrast to the MERS M^pro^-GC376 structure, the SARS-CoV-2 M^pro^ electron density indicated the formation of only one enantiomer for this inhibitor^12^. A strong hydrogen bond network is established from side chain of His163 and backbone amide of His164, and Glu166, with backbone contributions from Gly143, Ser144 and Cys145 defining the oxyanion hole (**Figure 3**). Together this provides strong binding and a low IC_50_ for the inhibitor. The glutamine surrogate in the substrate P1 position interacts with the side chain of His163, while the Leu in P2 inserts into a hydrophobic pocket, representing the S2 subsite of the enzyme. Similar to what was observed in the MERS-M^pro^-GC376 structure^12^, the benzyl ring and the β-lactam of the Gln surrogate forms a stacked hydrophobic interaction, which stabilizes the inhibitor in the active site of the protease. A close examination of the subsite for SARS-CoV-2 M^pro^ reveals regions to allow for future inhibitor development (**Extended data, Figure S6**).

**Figure 2.**
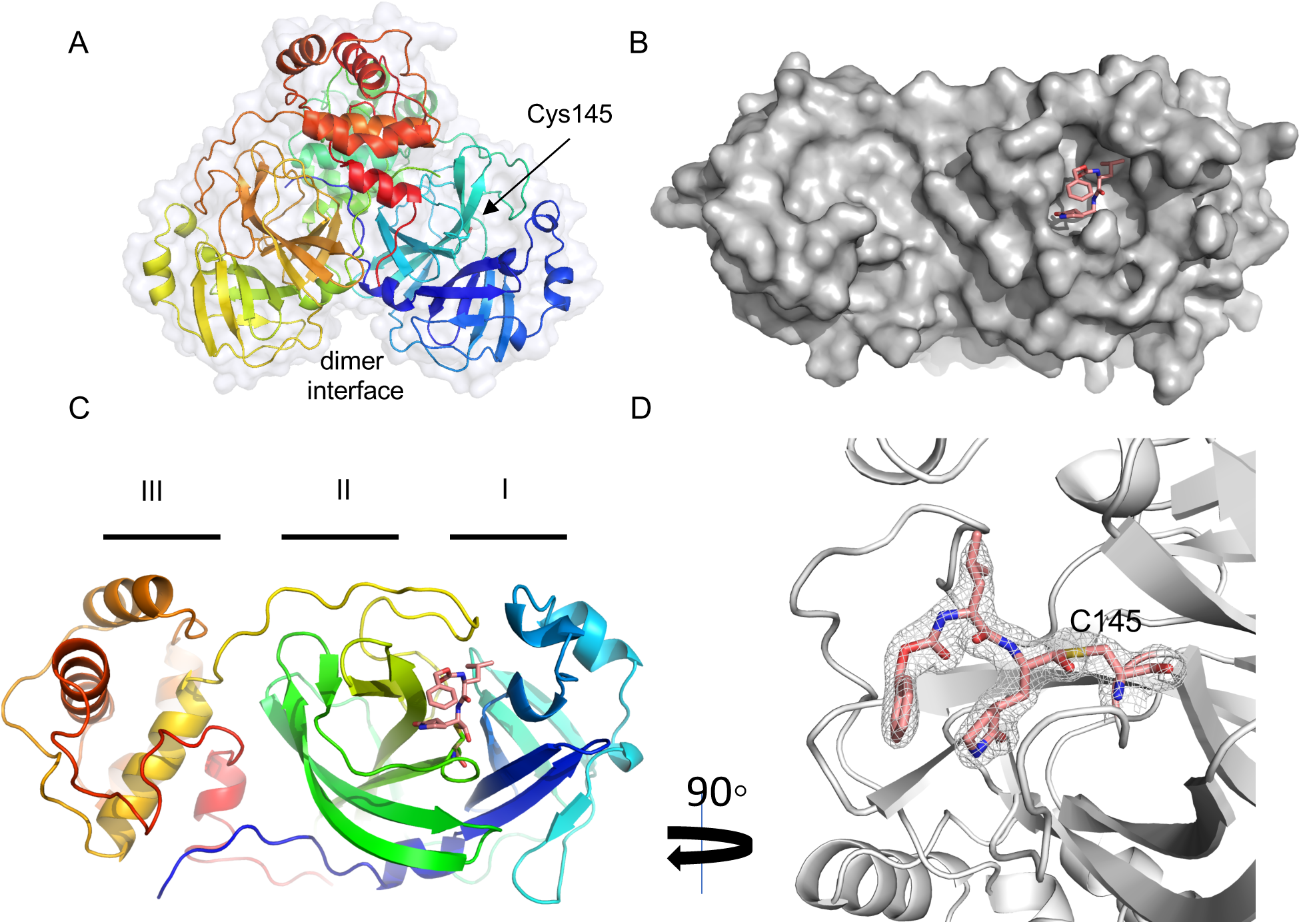
The crystal structure of SARS-CoV-2 M^pro^ in complex with GC373. **A)** Apo-SARS-CoV-2 M^pro^ forms a dimer (6WTM.pdb). **B)** Surface representation reveals the active site pocket in complex with GC376 (6WTJ.pdb). **C)** Ribbon representation of one SARS-CoV-2 protomer in complex with inhibitor GC376 binding in domain II. **D)** GC376 interacts covalently with the active site cysteine of SARS-CoV-2-M^pro^. Electron density if shown in grey mesh.

**Figure 3.**
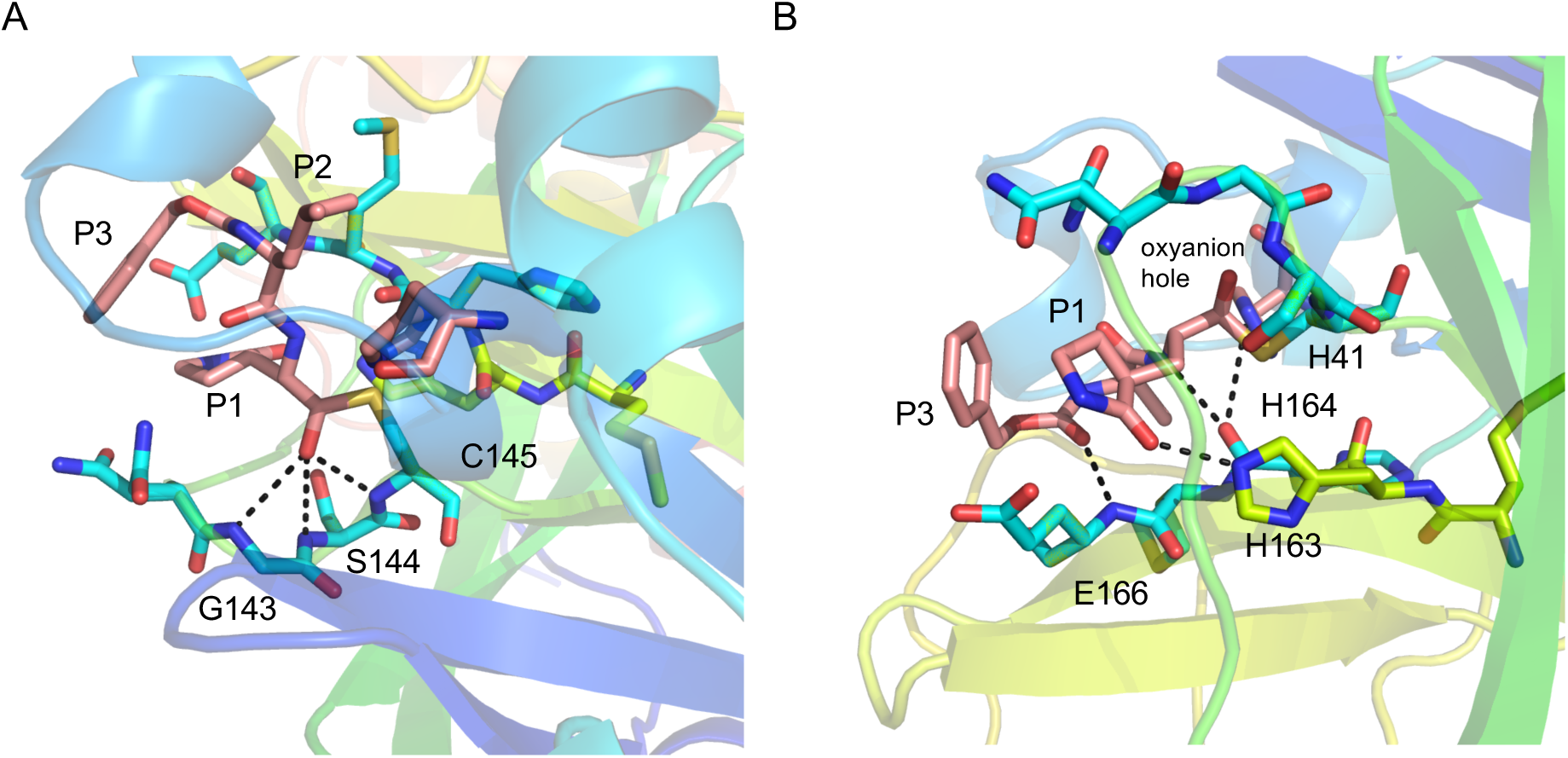
GC373 binds in the active site pocket of SARS-CoV-2 M^pro^. **A)** GC373 forms a covalent bond with Cys145, and the oxyanion is stabilized by backbone H-bonds with Gly143, Ser144, and Cys145. **B)** H-bonds are established with GC373 and His163 side chain as well as backbone of residues H164 and E166, which is supported by the backbone of His41. SARS-CoV-2 M^pro^ is represented in cartoon representation with the inhibitor in pink color. Inhibitor is colored in pink and interacting residues are colored in blue. P1, P2 and P3 of the peptidyl-inhibitor are indicated. PDB Code:6WTJ.

**Table 1.** Catalytic parameters of SARS-CoV and SARC-CoV-2 M^pro^ mediated cleavage of a FRET-peptide substrate. Catalytic parameters were determined for SARS-CoV M^pro^ and SARS-CoV-2 M^pro^ with the Abz-SVTLQSG-Y(NO2)-R substrate. Experiments were conducted in duplicate with an N=3. Values are represented as mean ±SEM.

| Protease | $K_{0.5}$<br>($\mu\text{M}$) | $k_{\text{cat}}$<br>($\text{min}^{-1}$) | $k_{\text{cat}}/K_{0.5}$<br>($\text{min}^{-1}\mu\text{M}^{-1}$) | Hill coefficient |
| --- | --- | --- | --- | --- |
| SARS-CoV-2 M<sup>pro</sup> | 70±10 | 135±6 | 1.8±0.4 | 2.2 |
| SARS-CoV M<sup>pro</sup> | 52±17 | 30±2 | 0.6±0.2 | 1.9 |

To confirm the formation of a covalent hemithioacetal as a single enantiomer in the active site, GC373 was prepared with ^13^C label (>99%) at the aldehyde carbon and mixed in 7.8-fold excess with the M^pro^ protease from SARS-CoV-2 in deuterated buffer. HSQC NMR analysis (700 MHz) showed appearance of a single crosspeak signal (one isomer only) for the hemithioacetal carbon at 76 ppm (^13^C) and 5.65 ppm (^1^H) in accordance with previous chemical shift reports for hemithioacetals^7^ (**Extended data, Figure S7**).

## SARS-CoV-2 M^pro^ has enhanced catalytic activity compared to SARS-CoV M^pro^

Recent crystal structural analysis reported differences in the residues residing between the dimer interface of SARS-CoV-2 M^pro^ when compared with the SARS-CoV M^pro^^13^. Previous mutagenesis studies, which altered residues at the dimer interface of SARS-CoV M^pro^, enhanced catalytic activity 3.6 fold^19^. In agreement with this, our analysis shows that the catalytic turnover rate for SARS-CoV-2-M^pro^ (135±6 min^-1^) is almost 5 times faster than SARS-CoV M^pro^ (30±2 min^-1^) with our substrate Abz-SVTLQSG-Tyr^NO2^R (**Table 1**). With this FRET substrate, we demonstrate a higher catalytic efficiency with SARS-CoV-2 M^pro^ (1.8±0.4 min^-1^µM^-1^) compared to SARS-CoV M^pro^ (0.6±0.2 min^-1^µM^-1^). This finding is in contrast to recent reports where no differences were observed in the catalytic efficiency between SARS-CoV M^pro^ and SARS-CoV-2 M^pro^ using a different substrate: 3011±294 s^-1^M^-1^ (0.18±0.02 min^-1^µM^-1^) and 3426±416.9 s^-1^M^-1^ (0.2±0.03 min^-1^µM^-1^), respectively)^13^. It remains to be determined whether this influences the virulence of SARS-CoV-2.

## GC373 and GC376 are potent inhibitors of SARS-CoV-2 in cell culture

To test the ability of GC373 and GC376 to inhibit SARS-CoV-2, plaque reduction assays were performed on infected Vero E6 cells in the absence or the presence of increasing concentrations of either GC373 (**Figure 4A**) or GC376 (**Figure 4B**) for 48 hours. The results were plotted as a percent inhibition of the number of plaque forming units per well. The EC_50_ for GC373 was 1.5 μM while the EC_50_ for GC376 was 0.92 μM. To examine cell cytotoxicity, ATP production was measured using the CellTiter-Glo assay on either Vero E6 cells or A549 cells incubated in the presence of the inhibitors for 24 hours. The CC_50_s of both GC373 and GC376 were greater than 200 μM. To further examine their antiviral activities, quantitative RT-PCR was performed on supernatants from cells untreated, and GC373 (**Figure 4C**) or GC376 (**Figure 4D**) treated cell cultures. It was observed that both GC373 and GC376 are potent inhibitors of SARS-CoV-2 decreasing viral titers 3 logs, compared with a 2 log decrease in recently published results using other aldehyde compounds^16^. These results indicate that both GC373 and GC376 are potent inhibitors of SARS-CoV-2 with a therapeutic index of >200.

**Figure 4.**
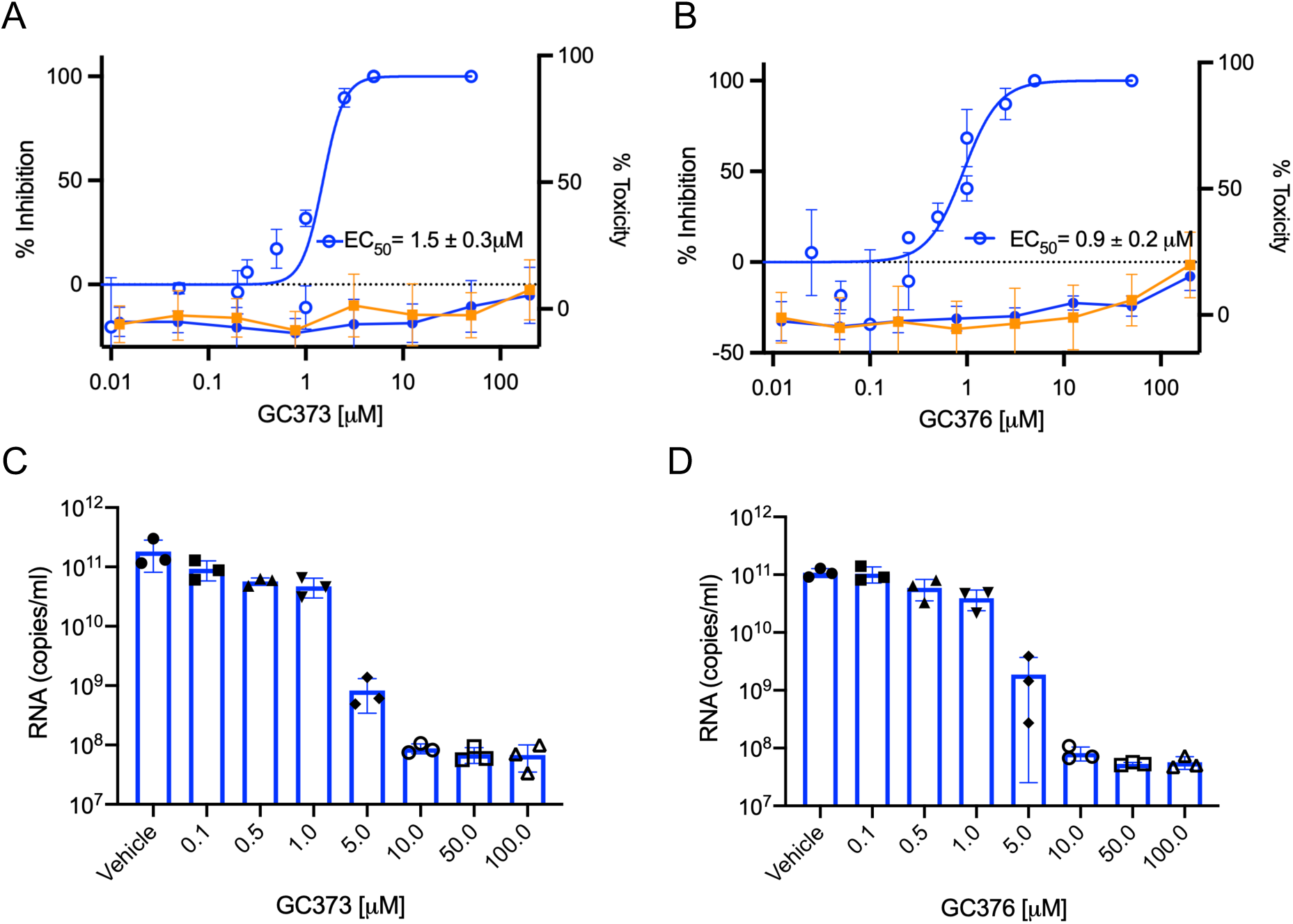
GC373 and GC376 potently inhibit SARS-CoV-2. **A)** and **B)** SARS-CoV-2 growth in Vero E6 cells was determined by plaque assays 48 h after infection in the presence of various concentrations of drug. Cytotoxicity was measured using the CellTiter-Glo assay. **A)** Percent inhibition of SARS-CoV-2 by GC373 in Vero E6 cells (blue open circles) and cytotoxicity in Vero E6 (blue closed circles) and A549 cells (orange). **B)** Percent inhibition of SARS-CoV-2 by GC376 in Vero E6 cells (blue open circles) and cytotoxicity in Vero E6 (blue, closed circles) and A549 cells (orange). **C)** and **D)** To measure the antiviral effect, Vero E6 cells were infected with MOI=0.01 of SARS-CoV-2 in triplicate without or with various concentration of GC373 **(C)** or GC376 **(D)** for 24 h and the supernatants harvested, RNA isolated and quantified by qRT-PCR. Values are mean ±SD.

Numerous drugs were designed originally to inhibit the SARS-CoV M^pro^ ^3^. However, the SARS outbreak of 2002 was controlled by public health measures and these compounds were never licensed. GC376 has been used to cure FIP in cats as the M^pro^ of FIP is effectively inhibited by its breakdown product GC373. Analogs of these drugs also inhibited the MERS CoV M^pro^ and blocked viral replication in cells at an EC_50_ of 0.5 µM^12^. Our studies show GC376 and GC373 to be effective inhibitors of SARS-CoV-2 M^pro^. Clearly these drugs need to be advanced quickly into human trials for COVID-19. SARS-CoV-2 is the cause of COVID-19 and is a virus with a significant mutation rate^20^. Also, in some patients the virus has persisted longer than 2 months with some possibility of re-infection^21^. Vaccines are critically important, but still likely a year or more away as this virus will likely present vaccine challenges.

There are many clinical trials testing drugs repurposed from their original indications. Remdesivir, a polymerase inhibitor developed as a treatment for Ebola virus^22^ is showing very promising early results and will likely be confirmed in clinical trials^23^. Another drug designed to inhibit RNA dependent RNA polymerase, including in coronaviruses, is EIDD 2801 (a N4-hydroxycytidine triphosphate that is incorporated into viral RNA to promote errors in progeny RNA). These examples of direct acting antivirals (DAAs) for COVID-19 are critically important^24^. Both GC373 and GC376 compounds are also DAAs designed specifically for coronaviruses. It is likely that several very potent drugs will be required to treat SARS-CoV-2 and to prevent the evolution of resistance they may need to be used in combination. We believe that GC373 and GC376 are candidate antivirals that should be accelerated into clinical trials for COVID-19.

## Online content

Any methods, additional references, Nature Research reporting summaries, source data, extended data, supplementary information, acknowledgements, peer review information; details of author contributions and competing interests; and statements of data and code availability are available online at:

## Author contributions

J.C.V., W.V. and T.L. contributed to inhibitor synthesis. W.V. contributed to FRET-substrate synthesis. M.J.v.B. contributed to cloning. E.A., M.J.v.B. and C.F. contributed to purified protein. C.F. and E.A. contributed to enzyme kinetics. M.J.L., H.S.Y. E.A., and M.B.K. contributed to crystallization and structure determination. W. V. and R. T. M. contributed to labelled NMR studies. D.L.T, M.A.J., H.A.S. and J.A.S contributed to viral inhibition and cell toxicity studies. M.J.L wrote the initial draft. All authors read and approved the manuscript.

## Acknowledgements

We would like to thank the staff at the Stanford Synchrotron Light Source, in particular Dr. Siliva Russi. M.J.L, J.C.V and D.L.T. acknowledge funding from CIHR and NSERC (COVID-19 SOF-549297-2019). D.L.T acknowledges support from Li Ka Shing Institute of Virology and the GSK Chair in Virology. W.V. was supported by an Alberta Innovates Graduate Scholarship and T.L. by a CIHR CGSM Scholarship.

## Extended data

**Figure S1.**
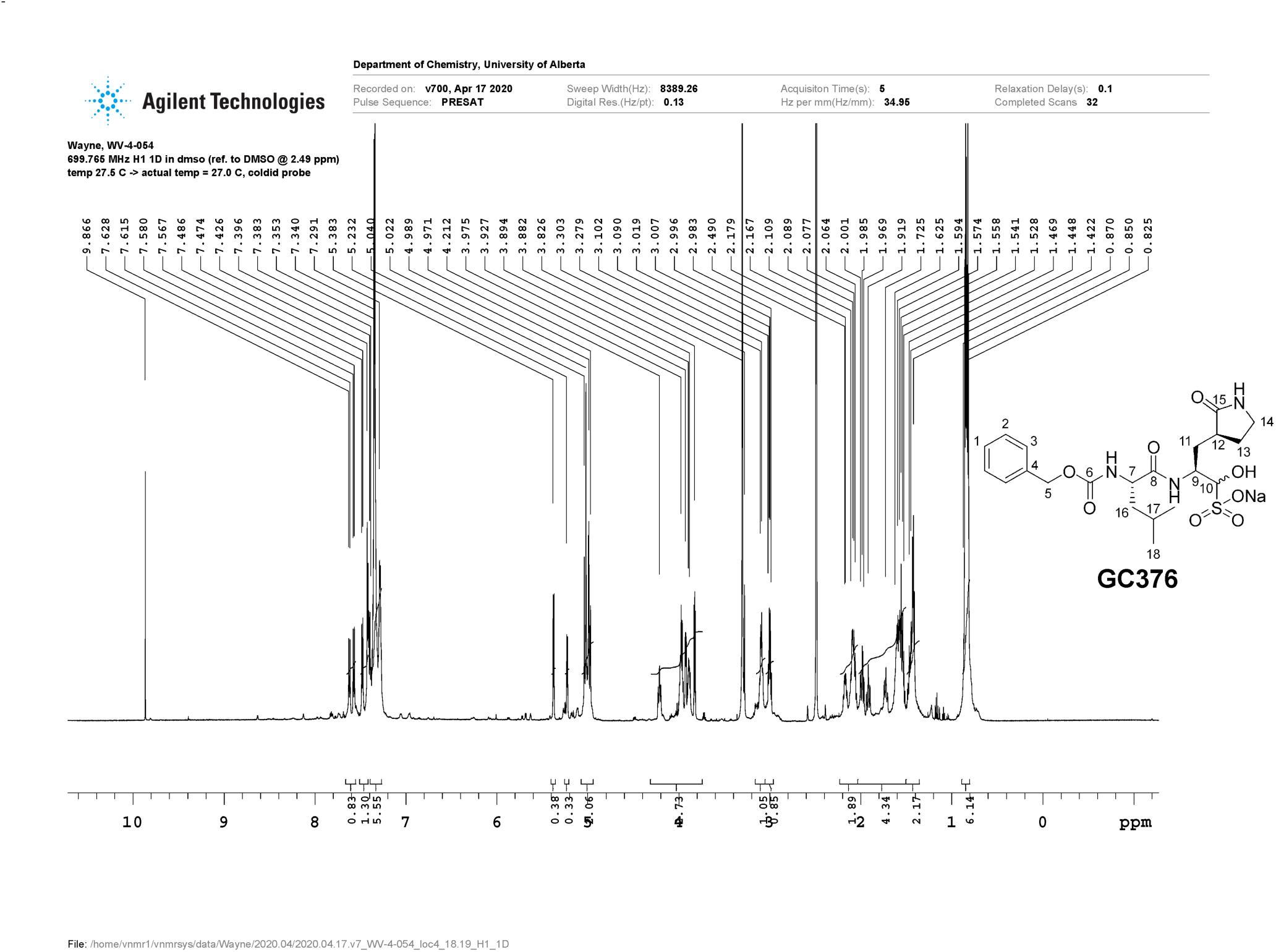
^1^H NMR spectrum of target inhibitor GC376.

**Figure S2.**
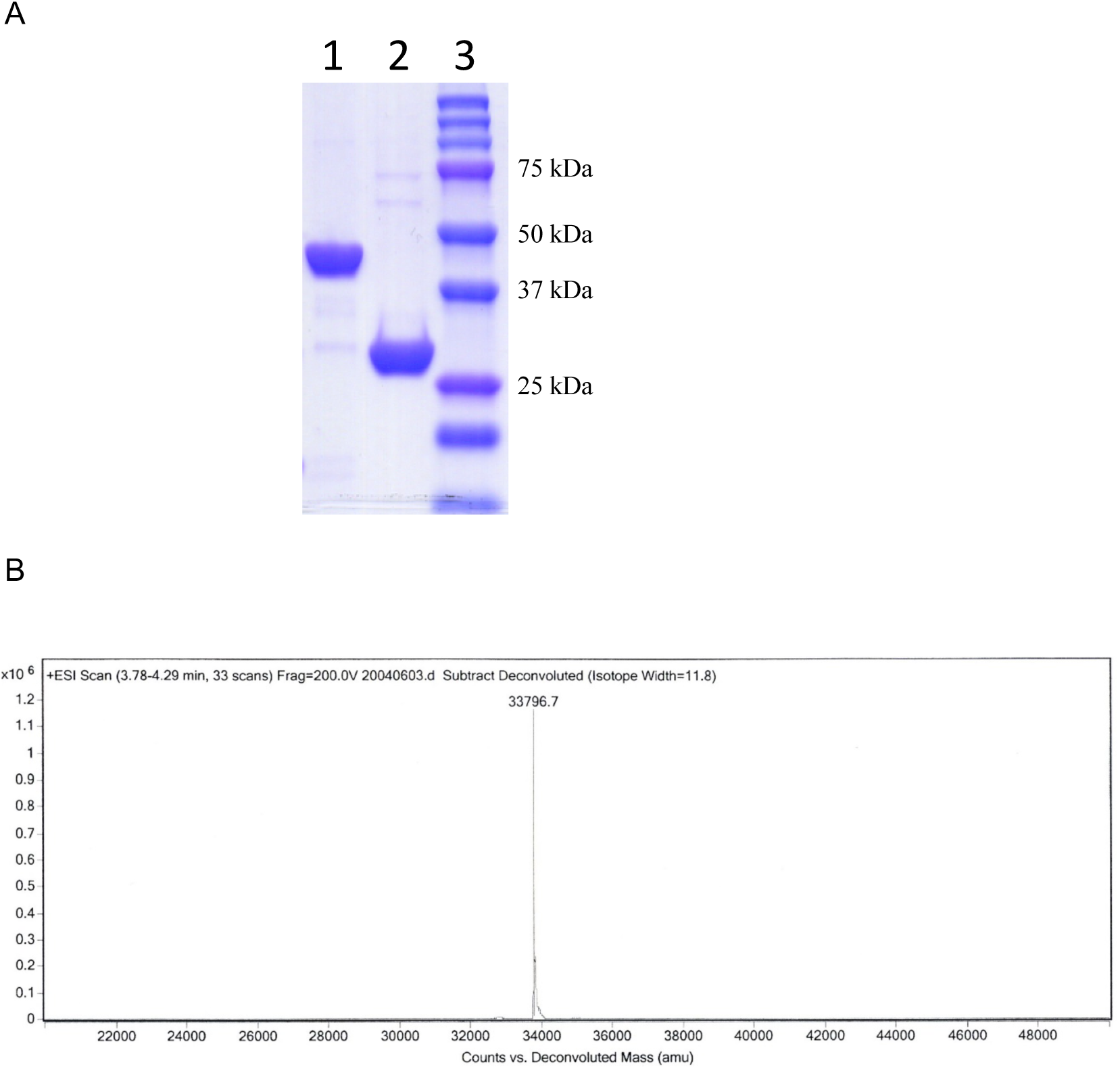
Purification of SARS-CoV-2 M^pro^. **A)** Coomassie stained SDS-PAGE of SARS-CoV-2 M^pro^ purification. Lane 1: SUMO-SARS-CoV-2 M^pro^ fusion protein; lane 2: SARS-CoV-2 M^pro^ after cleavage SUMO-tag; lane 3: protein MW ladder. **B)** ESI-MS of purified SARS-CoV-2 _M_pro.

**Figure S3.**
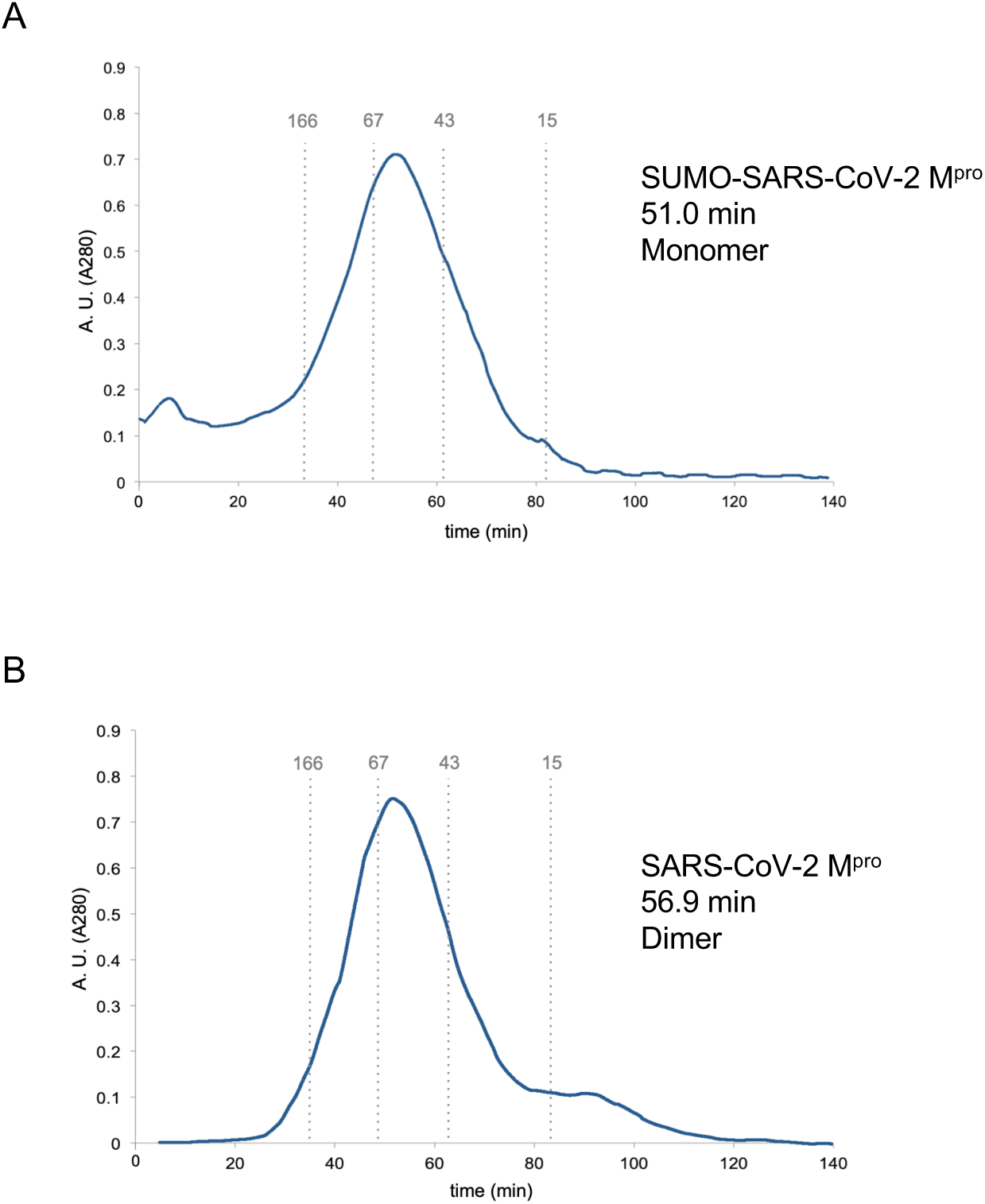
Gel filtration of SUMO-tagged SARS-CoV-2 M^pro^ and free SARS-CoV-2 M^pro^. **A)** The SUMO-tagged variant eluted as a monomeric species. **B)** The respective free SARS-CoV-2 M^pro^ exists predominantly as dimer (90% dimer). Retention times are provided at peaks. Calibration standards are indicated by grey lines and numbers represent kDa. AU= Absorbance at 280nm.

**Figure S4.**
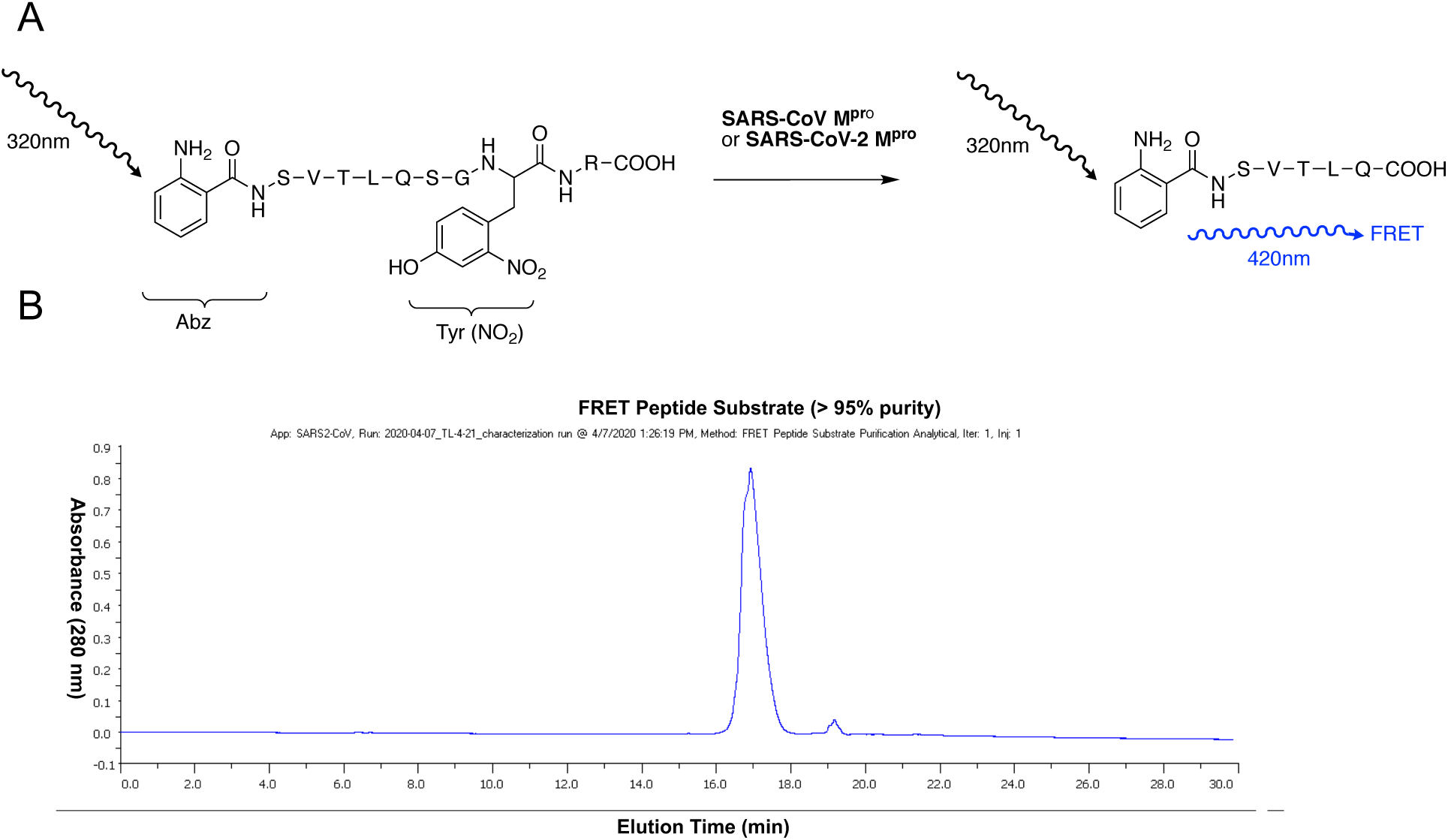
**A)** Molecular mechanism of the FRET assay involves a peptide 9-mer with fluorophore (Abz) and quencher (Tyr(NO_2_)). Upon cleavage by M^pro^ FRET emission at 420nm can be observed. **B)** HPLC chromatogram of FRET-substrate, Abz-SVTLQSG-Y(NO2)-R, indicating a purity >95%.

**Figure S5.**
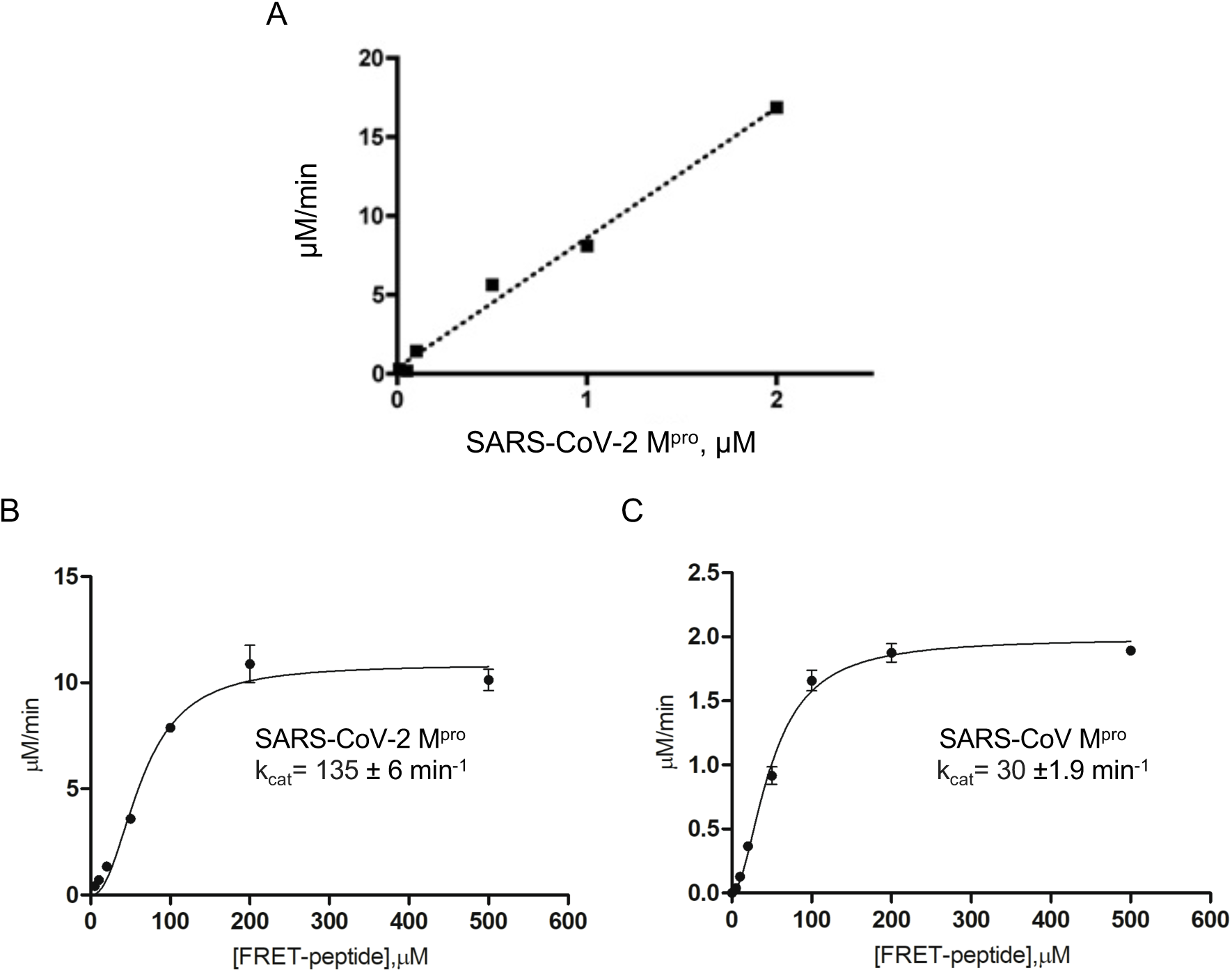
**A)** Linearity plot of enzyme activity. **B)** Representative Hill plots for Abz-SVTLQSG-Y(NO2)-R cleavage by SARS-CoV-2-M^pro^ and SARS M^pro^. N=3. Values represent mean±SEM.

**Figure S6.**
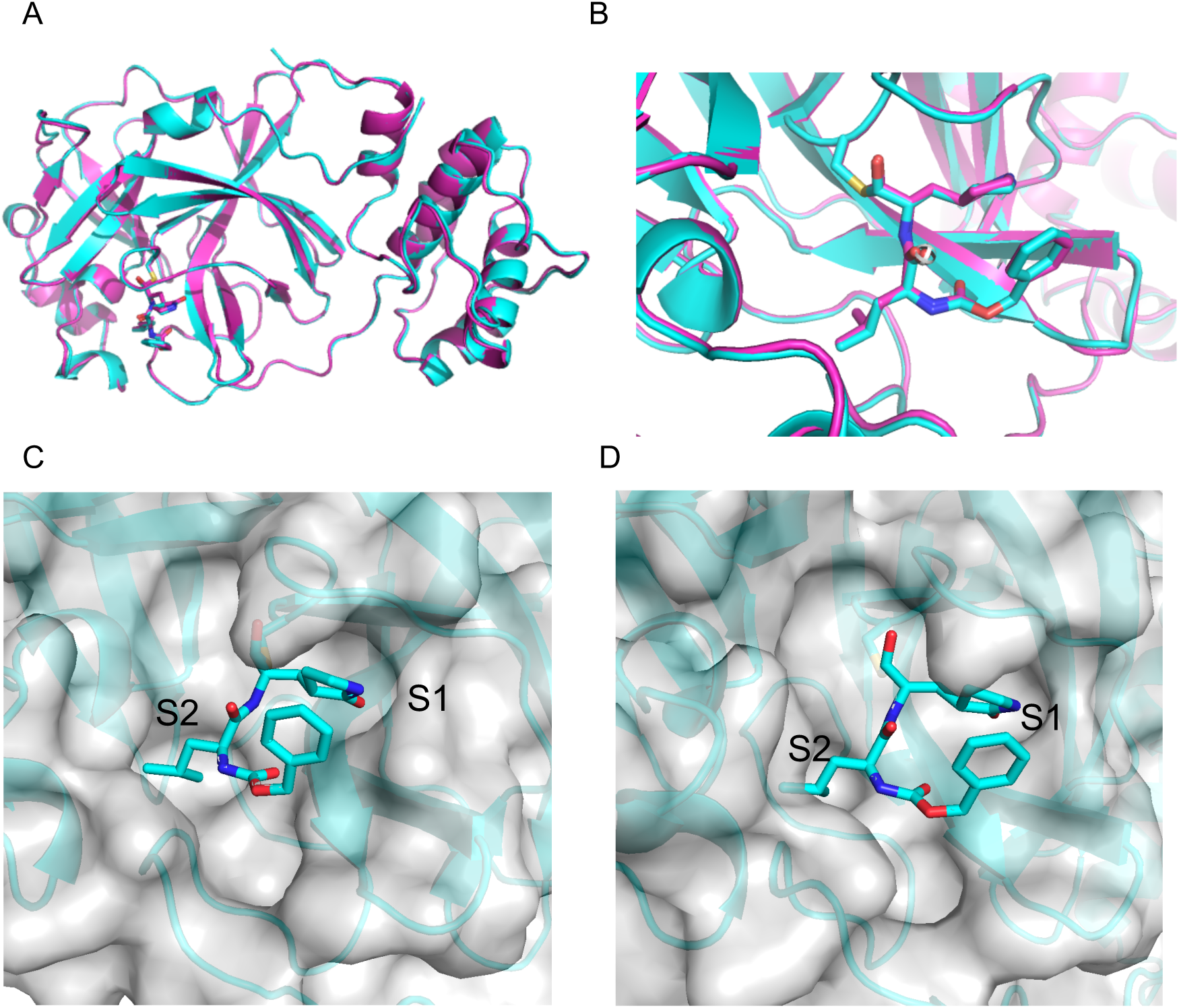
**A)** Alignment of crystal structure of SARS-CoV-2 M^pro^ soaked with GC373 (6WTK.pdb) or GC376 (6WTJ.pdb). GC373 is shown in pink, and GC376 in blue. **B)** Active site of SARS-CoV-2 M^pro^ with GC373 and GC376 show identical inhibitor structures, demonstrating the bisulphite adducts leave the GC376 compound to form GC373. Pink: GC376, blue=GC373. **C and D)** An examination of the fit in the enzyme subsites reveal regions to explore for novel inhibitor development.

**Figure S7.**
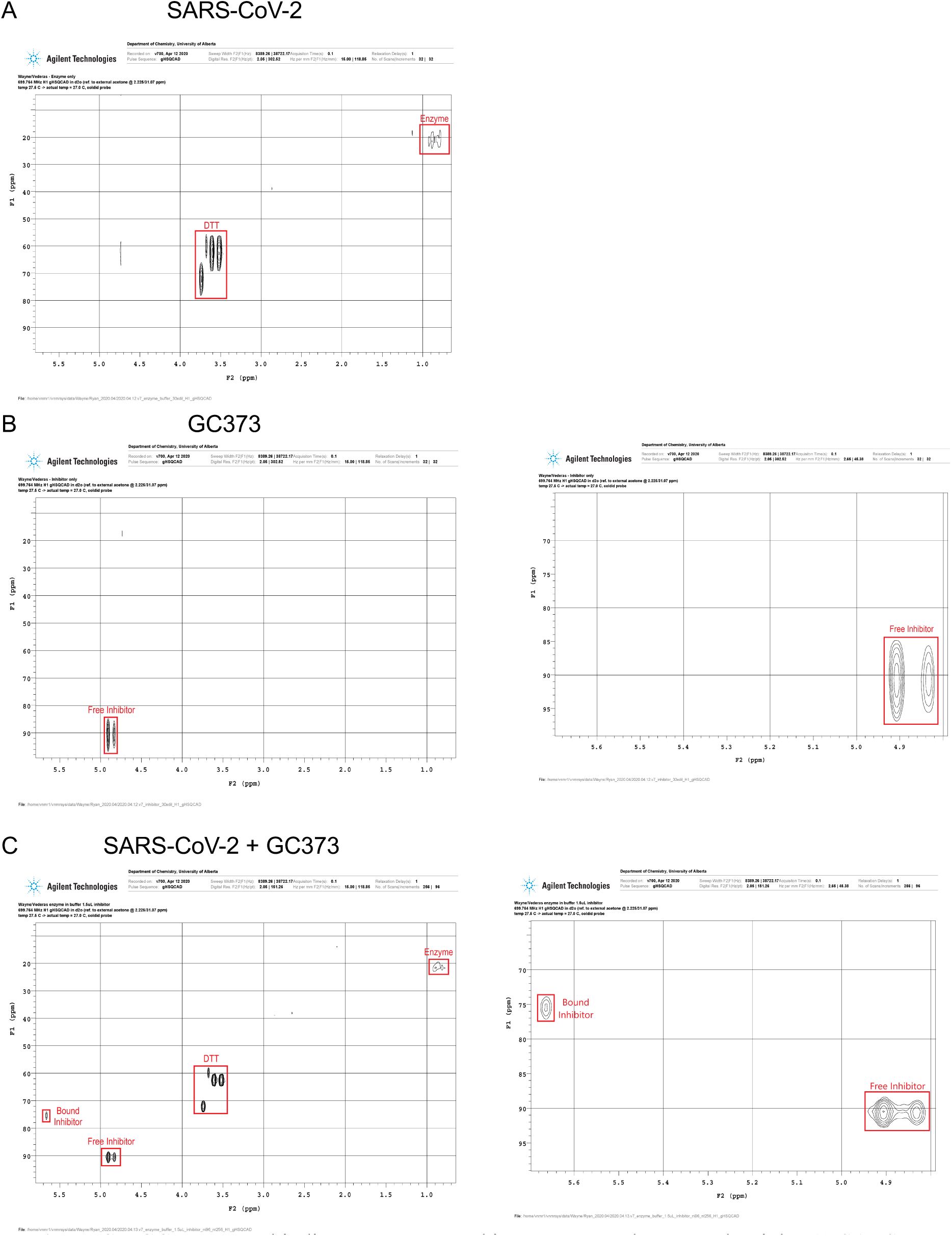
GC373 NMR binding assay. Zoomed in spectra are shown on the right. **A)** SARS-CoV-2 M^pro^ in the absence of GC373 inhibitor. **B)** GC373 inhibitor in the absence of SARS-CoV-2 M^pro^. **C)** Co-incubation of SARS-CoV-2 M^pro^ with GC373 inhibitor. A new crosspeak corresponding to the bound inhibitor can be observed, consistent with a bisulphite functionality.

**Table S1.** Diffraction data and model refinement statistics

| Protein/Ligand | SARS-CoV-2 Mpro | SARS-CoV2 Mpro-GC373 | SARS-CoV2 Mpro-GC376 |
| --- | --- | --- | --- |
| PDB Entry | 6WTM | 6WTK | 6WTJ |
| Data collection statistics |  |  |  |
| X-ray source | SSRL 12-2 | SSRL 12-2 | SSRL 12-2 |
| Wavelength [Å] | 0.97946 | 0.97946 | 0.97946 |
| V<sub>m</sub> [Å<sup>3</sup>/Da] | 2.0 | 1.97 | 2.04 |
| Solvent content [%] | 38.56 | 37.45 | 39.64 |
| Space group | P2<sub>1</sub> | C2 | C2 |
| Unit cell dimensions<br>a, b, c (Å)<br>α, β, γ (°) | 44.84 53.60 114.80 90<br>101.20 90. | 113.35 53.03 45.23 90<br>102.033 90 | 114.90 53.81 45.50<br>90 101.70 90 |
| Resolution range <sup>a</sup> [Å] | 3849-1.85 (1.89-1.85) | 34.04-2.0 (2.09-2.03) | 34.36-1.9 (1.95-1.90) |
| Number of observations<sup>a</sup> | 304078 (21067) | 115236 (7722) | 141537 (9860) |
| Number of unique reflections | 88303 (6137) | 34459 (2488) | 41270 (2925) |
| Completeness [%] <sup>a</sup> | 97.8 (92.2) | 96.0 (89.7) | 97.9 (93.4) |
| Mean I/σ(I) | 6.42 (1.0) | 5.60 (1.0) | 7.18 (1.0) |
| R-meas % | 15.1 | 16.2 | 11.8 |
| CC1/2 <sup>a</sup> | 99.6 (24) | 99.2(25) | 99.6 (25) |
| Refinement statistics |  |  |  |
| Number of unique reflections used for refinement | 88030 | 33622 | 41326 |
| R<sub>work</sub> /R<sub>free</sub> [%] | 20.80/25.18 | 20.13/25.99 | 20.67/25.47 |
| r.m.s.d. in bond lengths [Å]<br>bond angle (°) | 0.008/1.034 | 0.009/1.074 | 0.009/1.113 |
| Clashscore | 5.94 | 11.8 | 8.8 |
| Number of protein atoms | 4761 | 2367 | 2386 |
| Number of ligand atoms | N/A | 29 | 29 |
| Number of water molecules | 290 | 34 | 58 |
| Ramachandran plot |  |  |  |
| Preferred regions [%] | 95.01 | 97.70 | 95.68 |
| Allowed regions [%] | 4.49 | 1.32 | 3.32 |
| Outlier regions [%] | 0.50 | 0.99 | 1.33 |

<sup>a</sup> Highest resolution bin in parentheses

## Supporting Information

### Synthesis of GC376

#### General Characterization Methods

Nuclear magnetic resonance (NMR) spectra was obtained using an Agilent VNMRS 700 MHz of Agilent/Varian VNMRS 500 MHz spectrometer. For ^1^H (700 and 500 MHz) spectra, δ values were referenced to CDCl_3_ (7.26 ppm) or (CD_3_)_2_SO (2.50 ppm), and for ^13^C (175 of 125 MHz) spectra, δ values were referenced to CDCl_3_ (77.16 ppm), CD_3_OD or (CD_3_)_2_SO (39.52 ppm) as the solvents. Infrared spectras (IR) were recorded on a Nicolet Magna 750 or a 20SX FT-IR spectrometer. Cast film refers to the evaporation of a solution on an IR plate. Mass spectra were recorded on a ZabSpec IsoMass VG (high resolution electrospray ionization (ESI)). LC-MS analysis was performed on an Agilent Technologies 6220 orthogonal acceleration TOF instrument equipped with +ve and –ve ion ESI ionization, and full-scan MS (high-resolution analysis) with two-point lock mass correction operating mode. The instrument inlet was an Agilent Technologies 1200 SL HPLC system.

#### Reagents and Solvents

All commercially available reagents and protected amino acids were purchased and used without further purification unless otherwise noted. All the solvents used for reactions were used without further purification unless otherwise noted. Dry solvents refer to solvents freshly distilled over appropriate drying reagents prior to use.

Commercially available ACS grade solvents (> 99.0% purity) were used for column chromatography without any further purification.

#### Purificaton

All reactions and fractions from column chromatography were monitored by thin layer chromatography (TLC) using glass plates (5 × 2.5 cm) pre-coated (0.25 mm) with silica gel (normal SiO_2_, Merck 60 F254). Visualization of TLC plates was performed by UV fluorescence at 254 nm in addition to staining by KMnO_4_. Flash chromatography was performed using Merck type 60, 230-400 mesh silica gel at elevated pressures.

**Figure 1.**
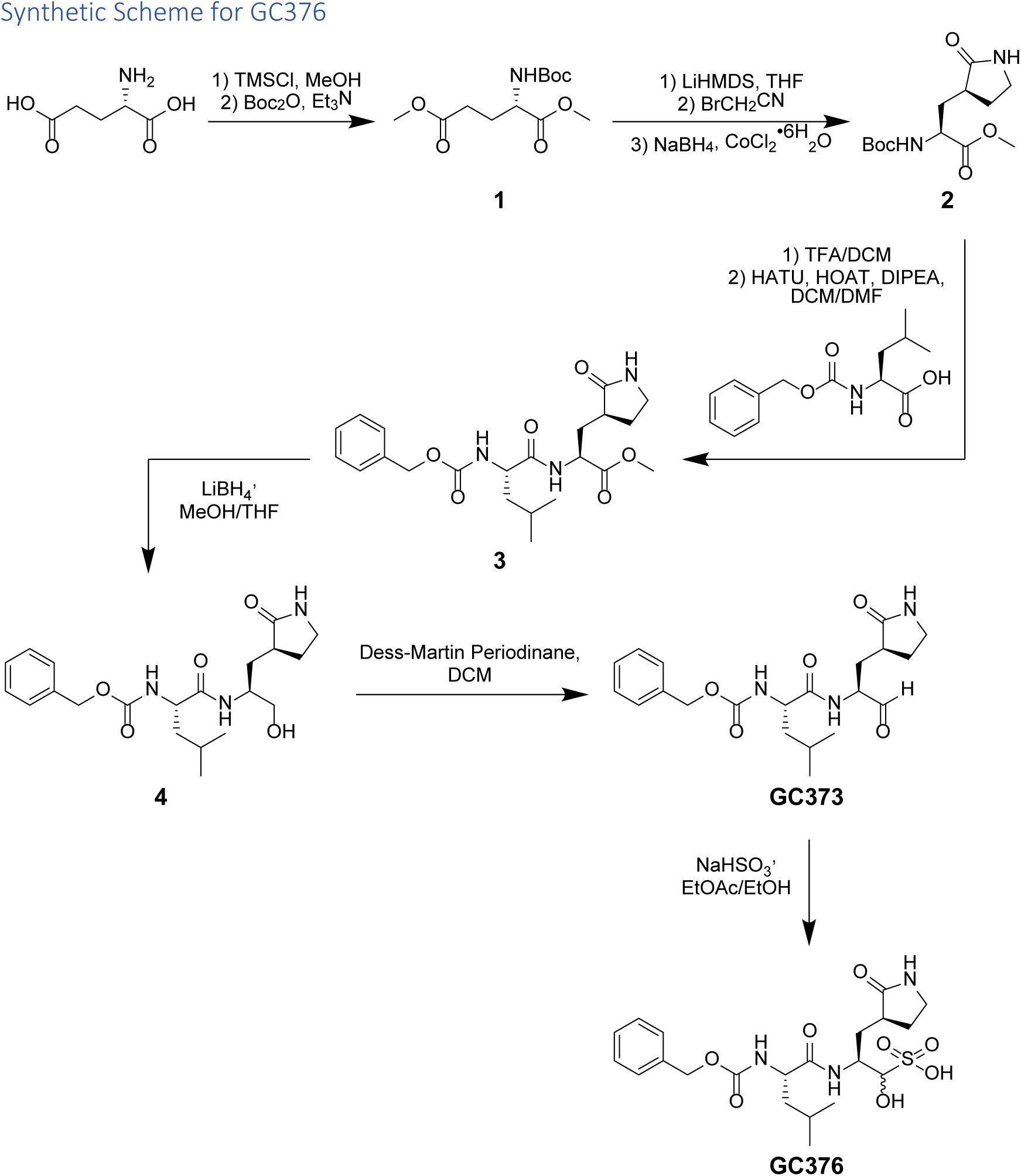
Synthetic scheme for the synthesis of GC376.

#### Dimethyl (tert-butoxycarbonyl)-L-glutamate (1)

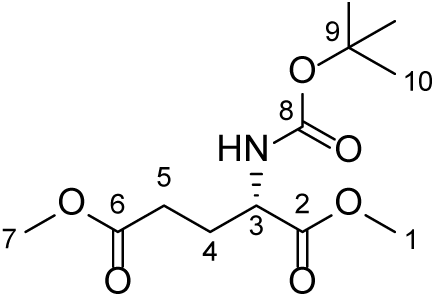

This known compound was synthesized based on a literature procedure^1^. *L*-Glutamic acid (30.00 g, 202.5 mmol, 1.0 equiv) was added to a flame-dried RBF under Ar, followed by the addition of dry MeOH (506.0 mL). The stirred suspension was then cooled to 0 °C, and TMSCl (113.1 mL, 891.7 mmol, 4.4 equiv) was added slowly over 15 minutes. The reaction mixture was stirred at 0 °C for 1.5 h, and then stirred overnight at rt. Triethylamine (182.5 mL, 1316 mmol, 6.5 equiv) was added to the reaction mixture slowly over 15 minutes. Next, Boc_2_O (48.62 g, 222.3 mmol, 1.1 equiv) was added in one portion, and then the reaction mixture was capped with pressure-equalizing Ar and stirred at rt for 3 h. The reaction mixture was then concentrated under reduced pressure to produce a white solid. The solid was triturated by suspension in Et_2_O (500 mL), and collected by filtration. The solid was washed with additional volumes of Et_2_O (3 × 250 mL), and then concentrated *in vacuo* to produce a crude oil. The crude product was purified using silica column chromatography (50% EtOAc in hexanes), and yielded a clear, yellow oil (50.44 g, 183.2 mmol, 91%): R*_f_ =* 0.61 on SiO_2_, 50% EtOAc in hexanes;

**IR** (DCM cast film, ν_max_ / cm^-1^) 3369, 2979, 2956, 1742, 1717, 1518, 1439, 1392, 1368, 1252, 1212, 1167

**^1^H NMR** (500 MHz, CDCl_3_) δ_H_ 5.13 (1H, d, *J* = 6.8 Hz, NH), 4.30 (1H, d, *J* = 5.0 Hz, H3)

3.71 (3H, s, H1), 3.65 (3H, s, H7), 2.31 (2H, m, H5), 2.15 (1H, app td, *J =* 13.2, 7.3, H4),

1.92 (1H, app dtd, *J =* 14.6, 8.3, 6.5, H4), 1.40 (9H, s, H10)

**^13^C NMR** (125 MHz, CDCl_3_) δ_C_ 173.3 (C6), 172.6 (C2), 155.3 (C8), 80.0 (C9), 52.8 (C3),

52.4 (C1), 51.7 (C7), 30.0 (C5), 28.3 (C10), 27.8 (C4)

**OR:** [α]_D_^26^ = 7.20 (*c* = 1.35, DCM)

**HRMS:** (ESI) Calcd for C_12_H_21_NNaO_6_ [M + Na]^+^ 298.1261, found 298.1260

#### (2S)-2-(tert-butoxycarbonylamino)-3-[(3’S)-2’-oxo-3’-pyrrolidinyl]propanoic acid methyl ester (2)

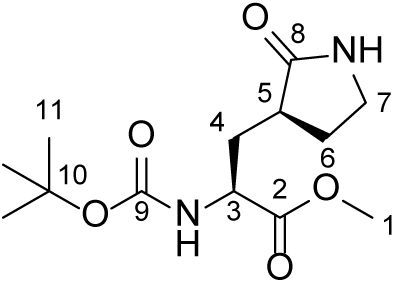

This known compound was synthesized based on a modified literature procedure^1^. **(1)** (1.00 g, 3.62 mmol, 1.00 equiv) was deposited in a flame-dried RBF under argon, to which 10.34 mL of freshly distilled THF was added. The oily starting material was dissolved at rt before LiHMDS (1.0 M in THF, 7.82 mL, 7.82 mmol, 2.16 equiv) was added over a period of 2 min at –78°C. The reaction mixture went from clear and nearly colourless to light yellow, and was allowed to stir at –78 °C for 1 h. Next BrCH_2_CN (0.270 mL, 3.88 mmol, 1.07 equiv) was slowly added over a period of 1 h (addition rate of ca. 0.02 mL / 5 min), while maintaining reaction temperature at –78 °C. The reaction mixture was then stirred at –78 °C over a period of 2 h. After 2 h, some starting material remained but the reaction mixture was quenched by addition of 5.2 mL of HCl at –78 °C. The reaction mixture was then removed from the cooling bath to allow the ice in the flask to melt, stirring for 50 min. The mixture was then extracted with EtOAc (3x). The combined EtOAc layers were washed with H_2_O (2x) and brine (1x). Drying over MgSO_4_ and concentration by rotovap furnishes a dark brown oil. This oil was then dissolved in 5.0 mL of DCM, to which activated charcoal (0.35 g) and silica (1.40 g) was added. The slurry was spun on a rotovap without heat or vacuum for 1 h. Afterwards, the charcoal and silica was filtered through a celite pad and the filtride washed with additional volumes of DCM. Concentration of the filtrate furnishes a yellow oil. This oil was then transferred to a flame-dried RBF under argon, to which CoCl_2_•6H_2_O (0.290 g, 1.22 mmol, 0.67 equiv) was added. The material was then dissolved in 20.0 mL of freshly distilled MeOH. The solution was then cooled to 0 °C and NaBH_4_ (0.462 g, 12.20 mmol, 6.74 equiv) was added in multiple portions over a period of 30 min. Upon addition of NaBH_4_, the reaction mixture immediately turned black and started bubbling. Once addition was finished and bubbling slowed, the reaction mixture was capped under a blanket of pressure-equalizing argon, removed from the ice bath, and allowed to warm to rt, stirring for 24 h. After 24 h, the reaction mixture was concentrated on the rotovap to a minimal volume. To the obtained residue was then added 1 M of citric acid at 0 °C. The mixture was further diluted with EtOAc and extracted with EtOAc (3x). The combined EtOAc layers were then washed with sat. NaHCO_3_ (2x) and brine (1x). Drying over Na_2_SO_4_ and removal of solvent furnishes a very pale yellow oil as crude. This material was then purified via flash column chromatography using an eluent system of EtOAc. Elution of product was monitored by KMnO_4_ staining (Rf = 0.16, EtOAc).

Concentration of product fractions and co-evaporation with Et_2_O furnishes the desired product as a white solid (0.280 g, 0.973 mmol, 54%).

**IR** (DCM cast film, ν_max_ / cm^-1^) 3290, 2977, 1745, 1698, 1523, 1440, 1392, 1367, 1276, 1213, 1168

**^1^H NMR** (500 MHz, CDCl_3_) δ_H_ 6.62 (1H, s, NH), 5.55 (1H, d, *J* = 7.8 Hz, NH), 4.29 (1H, m, H3), 3.72 (3H, s, H1), 3.36 – 3.31 (2H, m, H7), 2.48 – 2.40 (2H, m, H5, H6), 2.11

(1H, ddd, *J* = 14.3, 10.8, 3.7 Hz, H4), 1.88 – 1.40 (2H, m, H4, H6), 1.41 (9H, s, H11)

**^13^C NMR** (125 MHz, CDCl_3_) δ_C_ 179.8 (C8), 172.9 (C2), 155.8 (C9), 79.9 (C10), 52.4

(C1), 52.3 (C3), 40.4 (C7), 38.2 (C5), 34.1 (C4), 28.3 (C11), 28.1 (C6)

**OR:** [α]_D_^26^ = 3.65 (*c* = 0.57, DCM)

**HRMS:** (ESI) Calcd for C_13_H_22_N_2_NaO_5_ [M + Na]^+^ 309.1421, found 309.1423

#### Methyl (S)-2-((S)-2-(((benzyloxy)carbonyl)amino)-4-methylpentanamido)-3-((S)-2-oxopyrrolidin-3-yl)propanoate (3)

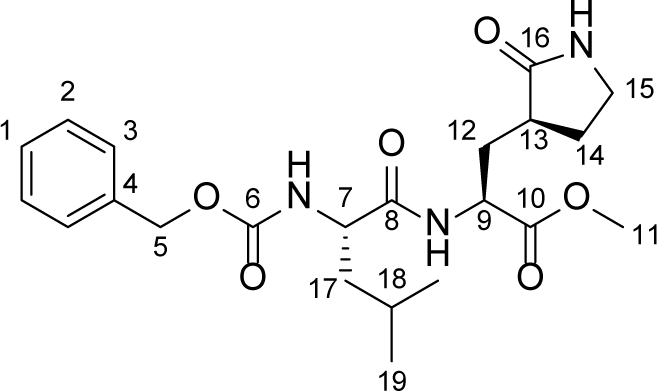

This known compound was synthesized *via* an alternative procedure. **(2)** (4.00 g, 13.97 mmol, 1.0 equiv) was dissolved in 140 mL of 50/50 TFA/DCM and stirred for 1 h at rt, with gas evolution being observed. The solution was then concentrated on rotovap and co-evaporated with DCM (5x). In a separate flame-dried RBF under argon was deposited Cbz-Leu-OH (90% purity, 4.12 g, 13.97 mmol, 1.0 equiv), HATU (5.31 g, 13.97 mmol, 1.0 equiv). The material was then dissolved in 70 mL of DMF. Next, HOAT (0.6 M in DMF, 2.33 mL, 1.40 mmol, 0.1 equiv) was added, followed by DIPEA (7.30 mL, 41.91 mmol, 3.0 equiv). The reaction mixture turned bright yellow and was incubated for 5 min. The previously concentrated Boc-deprotected material was then dissolved in 70 mL of DCM and added dropwise to the incubating solution, with the bright yellow colour quickly fading. The reaction mixture was the capped under a blanket of argon and allowed to react at rt for 1.5 h. Next, the reaction mixture was diluted with H_2_O and EtOAc. The layers were separated and the H_2_O layer was further extracted with EtOAc (3x). The combined EtOAc layers were then washed with sat. NaHCO_3_ (2x), 1 M HCl (2x), and brine (1x). Drying over Na_2_SO_4_ and concentrating on the rotovap furnishes a crude yellow oil. This material was used without further purification but may be purified by flash column chromatography over silica, using an eluent system of pure EtOAc. Product elution was monitored by TLC and KMnO_4_ staining (R_f_ = 0.43, 5/95 MeOH/EtOAc), and concentration of product fractions furnishes a transparent, slightly yellow oil as the desired product (6.056 g, 13.97 mmol, 100%).

**IR** (DCM cast film, ν_max_ / cm^-1^) 3275, 3063, 2955, 2871, 1688, 1538, 1455, 1439, 1385, 1268, 1176, 1134

**^1^H NMR** (700 MHz, CDCl_3_) δ_H_ 7.84 (1H, d, *J* = 5.3 Hz, NH), 7.37—7.27 (5H, m, H1, H2, H3), 5.97 (1H, s, NH), 5.34 (1H, d, *J* = 7.0 Hz), 5.09 (2H, s, H5), 4.51—4.42 (1H, m,

H9), 4.36—4.27 (1H, m, H7), 3.72 (3H, s, H11), 3.35—3.25 (2H, m, H15), 2.42—2.32

(2H, m, H13, H14), 2.21—2.10 (1H, m, H12), 1.95—1.88 (1H, m, H12), 1.86—1.78 (1H, m, H14), 1.78—1.71 (1H, m, H18), 1.71—1.65 (1H, m, H17), 1.56—1.47 (1H, m H17), 0.99—0.91 (6H, m, H19)

**^13^C NMR** (175 MHz, CDCl_3_) δ_C_ 179.7 (C16), 172.8 (C8), 172.1 (C10), 156.1 (C6), 136.4

(C4), 128.5 (C2), 128.1 (C3), 128.0 (C1), 66.9 (C5), 53.4 (C7), 52.4 (C11), 51.7 (C9),

42.3 (C17), 40.5 (C15), 38.4 (C13), 32.9 (C12), 28.6 (C14), 24.6 (C18), 22.9 (H19), 22.1 (H19)

**OR:** [α]_D_^26^ = −12.51 (*c* = 0.78, DCM)

**HRMS:** (ESI) (ESI) Calcd for C_22_H_31_N_3_NaO_6_ [M + Na]^+^ 456.2105, found 456.2101

#### N^2^-[(Benzyloxy)carbonyl]-N-{(2S)-1-hydroxy-3-[(3S)-2-oxo-3-pyrrolidinyl]-2-propanyl}-L-leucinamide (4)

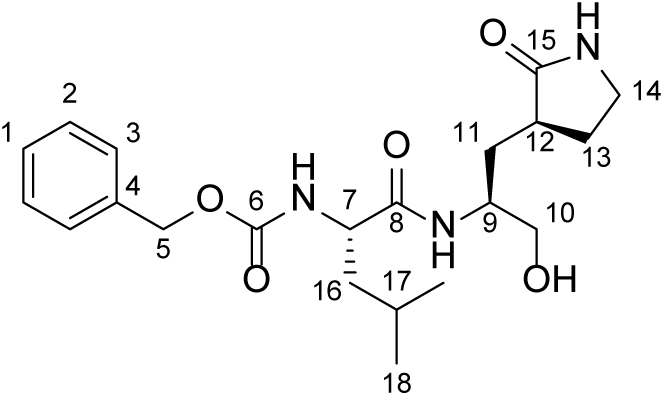

This compound was synthesized based on a literature procedure^2^. **(3)** (0.167 g, 0.385 mmol, 1.0 equiv) was deposited in a flame-dried round bottom flask under argon and dissolved in 2.31 mL of dry THF. To the RM was then added LiBH_4_ (2.0 M in THF, 0.578 mL, 1.156 mmol, 3.0 equiv) dropwise. Addition causes RM to turn bright yellow in colour. Continued addition eventually results in fading of yellow colour. Gas evolution occurred. After gas evolution ceased, 1.16 mL of dry MeOH was added dropwise. This was followed by a second gas evolution event. RM was allowed to stir at rt for 1.5 h under a blanket of argon. Reaction mixture was worked up by quenching with 1 M HCl until pH 1—2. The reaction mixture was then concentrated on rotovap and the resulting residue was dissolved in EtOAc and brine. The layers were separated and the EtOAc layer was dried over Na_2_SO_4_. Concentration and co-evaporation with Et_2_O furnishes a white foam as the product (R_f_ = 0.14, 5/95 MeOH/EtOAc) (0.14 g, 0.345 mmol, 90%) with no further purification required.

**IR** (DCM cast film, ν_max_ / cm^-1^) 3282, 3065, 2955, 2871, 1686, 1539, 1456, 1442, 1386, 1369, 1263, 1173

**^1^H NMR** (500 MHz, CDCl_3_) δ_H_ 7.79—7.63 (1H, m, NH), 7.40—7.27 (5H, m, H1, H2, H3),

6.51—6.00 (1H, m, NH), 5.74—5.41 (1H, m, NH), 5.09 (2H, s, H5), 4.32—4.16 (1H, s,

H7), 4.04—3.90 (1H, s, H9), 3.73—3.40 (3H, m, H10, OH), 3.34—3.18 (2H, m, H14),

2.55—2.25 (2H, m, H12, H13), 2.06—1.94 (1H, m, H11), 1.85—1.73 (1H, m, H13),

1.73—1.56 (3H, m, H11, H16, H17), 1.58—1.44 (1H, m, H16), 1.01—0.84 (6H, m, H18)

**^13^C NMR** (125 MHz, CDCl_3_) δ_C_ 181.0 (C15), 173.7(C8), 156.2 (C6), 136.4 (C4), 128.5

(C2), 128.1 (C3), 128.0 (C1), 66.9 (C5), 66.1 (C10), 53.9 (C7), 51.3 (C9), 42.3 (C16),

40.6 (C14), 38.6 (C12), 32.1 (C11), 28.9 (C13), 24.8 (C17), 23.0 (C18), 22.0 (C18)

**OR:** [α]_D_^26^ = −21.20 (*c* = 1.05, DCM)

**HRMS:** (ESI) Calcd for C_21_H_32_N_3_O_5_ [M + H]^+^ 406.2336, found 405.2264

#### (2S)-2-((S)-2-(((benzyloxy)carbonyl)amino)-4-methylpentanamido)-3-(2-oxopyrrolidin-3-yl)propanal (GC373)

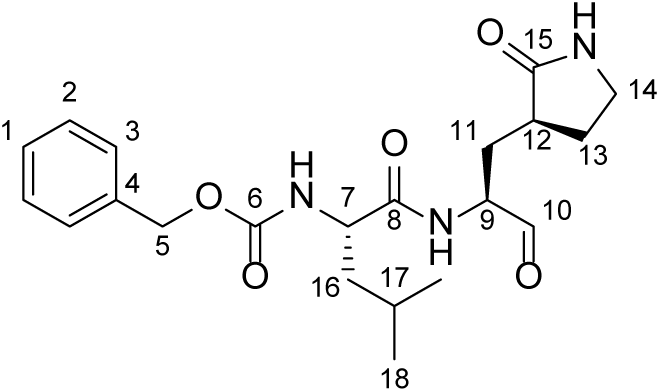

This compound was synthesized based on a literature procedure^2^. **(4)** (0.319 g, 0.786 mmol, 1.0 equiv) was dissolved in dry DCM under argon. Dess—Martin periodinane (0.500 g, 1.179 mmol, 1.5 equiv) was then added at 0 °C. The reaction mixture was capped under a blanket of argon and allowed to warm to rt. The reaction was then allowed to stir at rt for 3h. Reaction mixture was then quenched with the addition of 3 mL of 10% Na_2_S_2_O_3_ followed with stirring for 15 min. The layers were then separated and the DCM layer was washed sequentially with 10% Na_2_S_2_O_3_ (1 x 3 mL), sat.

NaHCO_3_ (2 x 3 mL), H2O (2 x 3 mL), and brine (2 x 3 mL). The DCM layer was then dried over Na_2_SO_4_ and concentrated to furnish a yellow residue. The TLC of this crude is very messy (extensive streaking) but the product was successfully purified *via* flash column chromatography on silica using an eluent system of 2.5/97.5 MeOH/EtOAc.

Elution of product was monitored by TLC and KMnO_4_ staining (R_f_ = 0.14, 2.5/97.5 MeOH/EtOAc). Concentration of product fractions furnishes an oil that solidifies to a yellow foam upon co-evaporation with Et_2_O (0.206 g, 0.511 mmol, 65%).

**IR** (DCM cast film, ν_max_ / cm^-1^) 3287, 3064, 2957, 2871, 1691, 1535, 1456, 1440, 1386, 1369, 1265, 1119

Please note that this compound exists as a mixture of diastereomers due to rapid epimerization at the alpha carbon of alpha-amido aldehydes. Although the ^1^H NMR spectra does not appear to readily show this, some ^13^C NMR signals do appear separated and have been noted as pairs where appropriate.

**^1^H NMR** (500 MHz, CDCl_3_) δ_H_ 9.51 (1H, app d, *J* = 33.9 Hz, H10 of diastereomers), 8.42 (1H, app dd, *J* = 77.0, 4.62 Hz, NH of diastereomers), 7.42—7.27 (5H, m, H1, H2, H3), 6.18 (1H, app d, *J* = 77.0 Hz, NH of diastereomers), 5.45 (1H, app dd, *J* = 36.5, 8.25 Hz, NH of diastereomers), 5.16—5.00 (2H, d, *J* = 9.5 Hz, H5), 4.55—4.07 (2H, m, H7, H9), 3.40—3.17 (2H, m, H14), 2.58—2.20 (2H, m, H12, H13), 2.14—2.00 (1H, m, H11), 2.00—1.88 (1H, m, H11), 1.88—1.80 (1H, m, H13), 1.79—1.62 (2H, m, H16, H17), 1.62—1.44 (1H, m, H16), 1.03—0.84 (6H, m, H18)

**^13^C NMR** (125 MHz, CDCl_3_) δ_C_ 200.8/199.7 (C10), 180.2/180.1 (C15), 173.7/173.4 (C8),

156.1 (C6), 136.4 (C4), 128.6/128.5 (C2), 128.2/128.1 (C3), 128.0 (C1), 67.0/66.9 (C5),

57.9/57.0 (C7), 53.7/53.6 (C9), 42.3/41.9 (C16), 40.6 (C14), 38.2/37.7 (C12), 29.7/29.1

(C11), 28.8/28.3 (C13), 24.8 (C17), 23.0/23.0 (C18), 22.0/21.9 (C18)

**OR:** [α]_D_^26^ = −2.93 (*c* = 0.43, DCM)

**HRMS:** (ESI) Calcd for C_21_H_30_N_3_O_5_ [M + H]^+^ 404.2180, found 404.2173

#### Sodium (2S)-2-((S)-2-(((benzyloxy)carbonyl)amino)-4-methylpentanamido)-1-hydroxy-3-(2-oxopyrrolidin-3-yl)propane-1-sulfonate (GC376)

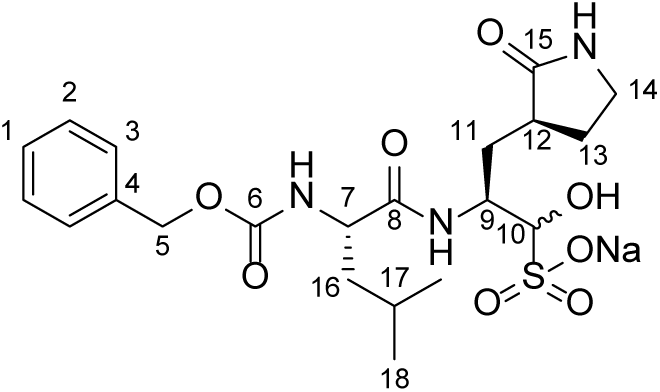

This compound was synthesized based on a literature procedure^2^. **GC373** (0.030 g, 0.074 mmol, 1.0 equiv) was deposited in a vial and dissolved in 0.296 mL of dry EtOAc and 0.178 mL of absolute EtOH. To this was added 1 M NaHSO_3_ (0.074 mL, 0.074 mmol, 1.0 equiv). The reaction vessel was then capped and stirred at 50 °C for 3 h. Afterwards, the mixture was filtered to remove solids and the solids were thoroughly washed with additional volumes of EtOH. The combined washings were dried over Na_2_SO_4_ and filtered again. Concentration of the filtrate furnishes a yellow oil. To this oil was added 0.5 mL of Et_2_O, causing a white solid to crash out after mixing. The mixture was then centrifuged and the Et_2_O was removed. This step was repeated once more. The resultant white solid was then treated with Et_2_O (0.444 mL) and EtOAc (0.222 mL). Mixing for 5 min followed by centrifuge and removal of solvent furnishes the white bisulfite adduct as the product. The material was dried on hi-vac overnight to remove residual solvent (0.018 g, 0.037 mmol, 50%). A check by LCMS shows the desired adduct as the major product (0.255 g, 0.050 mmol, 68%) and was used without further purification.

**IR** (H_2_O cast film, ν_max_ / cm^-1^) 3291, 3090, 3066, 3035, 2957, 2872, 1674, 1527, 1455, 1387, 1368, 1216

Please note that this compound exists as a mixture of diastereomers due to rapid epimerization at the alpha carbon of alpha-amido aldehydes. Although the ^1^H NMR spectra does not appear to readily show this, some ^13^C NMR signals do appear separated and have been noted as pairs where appropriate.

**^1^H NMR** (500 MHz, DMSO-d6) δ_H_ 7.65—7.56 (1H, m NH), 7.50—7.42 (1H, m NH),

7.39—7.31 (5H, m, H1, H2, H3), 5.38 (1H, d, *J* = 4.5 Hz, NH), 5.24 (1H, d, *J* = 4.5 Hz,

OH), 5.07—4.96 (2H, m, H5), 4.25—3.79 (3H, m, H7, H9, H10), 3.14—3.07 (1H, m,

H14), 3.04—2.98 (1H, m, H14), 2.22—2.04 (2H, m, H12, H13), 2.03—1.52 (4H, m, H11,

H13, H17), 1.50—1.40 (2H, m, H16), 0.85 (6H, ddd, *J* = 12.4, 8.9, 4.4 Hz, H18)

**^13^C NMR** (125 MHz, DMSO-d6) δ_C_ 179.0 (C15), 171.8 (C8), 156.0 (C6), 137.1 (C4),

128.2/128.2 (C2), 127.6 (C3), 127.6/127.6 (C1), 84.4/83.7 (C10), 65.3 (C5), 53.6/53.5

(C9), 49.1/48.6 (C7), 40.5 (C16), 40.7/40.4 (C14), 37.8/37.7 (C12), 31.8 (C11),

27.5/27.3 (C13), 24.2/24.2 (C17), 23.1/23.0 (C18), 21.4/21.3 (C18)

**OR:** [α]_D_^26^ = −37.38 (*c* = 0.31, H_2_O)

**HRMS:** (ESI) Calcd for C_21_H_32_N_3_O_8_S [M + H]^+^ 486.1905, found 486.1896

#### Synthesis of ^13^C Labelled GC373

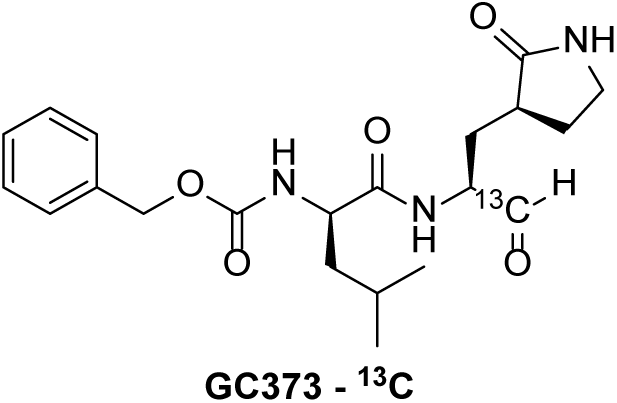

Synthesis of ^13^C labelled GC373 was carried out *via* the above documented methods. All characterization data was in agreement as described above.

**HRMS:** (ESI) Calcd for C_20_^13^CH_30_N_3_O_5_ [M + H]^+^ 405.2214, found 405.2215

### FRET Substrate Peptide Synthesis

#### General 2-Chlorotrityl Chloride Resin Loading Procedure

2-Chlorotrityl chloride resin was transferred to a SPPS vessel and washed with dry CH_2_Cl_2_ (2 × 10 mL) and then dry DMF (2 × 10 mL) for one min each, and then bubbled under Ar in dry DMF (10 mL) for 10 min. The desired Fmoc-protected amino acid (1.0 equiv, based on desired resin loading) and DIPEA (5.0 equiv) were suspended in 10 mL of a 50/50 mixture of dry CH_2_Cl_2_/DMF. This solution was bubbled under Ar for 2.5 h to load the desired amino acid onto the solid support, continually topping up the CH_2_Cl_2_ to maintain an approximately 10 mL volume. To end cap any remaining trityl groups, dry MeOH was added to the vessel (0.8 mL per gram of resin) and bubbled under Ar for 15 minutes. After draining, the resin was washed with dry DMF (3 × 10 mL), dry CH_2_Cl_2_ (3 × 10 mL), and then with dry MeOH (3 × 10 mL) for one min each. The resin was dried thoroughly and then stored at –20 °C under Ar.

#### General Automated SPPS Elongation Method

All peptides were synthesized on a PreludeX (Gyros protein technologies). SPPS was carried out on a 0.1 mmol scale using Fmoc chemistry on 2-chlorotrityl resin (0.8 mmol/g). Commercially available Fmoc-protected amino acids were loaded on the peptide synthesizer as 0.2 M solutions in DMF. All amino acids were coupled using 1-[Bis(dimethylamino)-methylene]-1H-1,2,3-triazolo[4,5-b]pyridinium3-oxid hexafluoro-phosphate (HATU) as the activating agent with a coupling time of 1 h. Fmoc residues were deprotected using a 20% solution of piperidine in DMF.

#### General Method for Cleavage of Peptide from Resin

Resin-bound analogue was suspended in 95/2.5/2.5 TFA/TIPS/H_2_O with shaking for 2-3 h. The resin was removed *via* filtration through glass wool, rinsed with TFA, and the solution concentrated *in vacuo*. Cold diethyl ether (2 × 5 mL) was added to triturate the crude residue. The diethyl ether was decanted and briefly centrifuged for 3 minutes at 13000 rpm to pellet any residual peptide. The ether was removed and the peptide pellet was then dried thoroughly by centrifugation in a vacuum centrifuge for 5 minutes. The pellet and triturated crude residue were pooled together and dissolved in 0.1% aqueous TFA.

#### HPLC Purification Method

The FRET peptide substrate was purified using a C_18_ RP-HPLC column with aqueous 0.1% TFA (solvent A) and 0.1% TFA in acetonitrile (solvent B) as eluents. The analytical purification method used was: 0 – 3 min 10% B, 3 – 4.5 min 10% – 25% B, 4.5 – 14.5 min 25% – 40% B, 14.5 – 17 min 40% – 90% B, 17 – 19.5 min 95% B, 19.5 – 20.5 min 95% – 10% B, 20.5 – 30 min 10% B.

#### Synthesis of FRET Peptide Substrate

Fmoc-L-Arg(Pmc)-OH was loaded onto 2-chlorotrityl chloride resin (0.18 mmol/g) using the aforementioned general procedure. The resin was elongated using automated SPPS, introducing amino acids in the following order: Fmoc-Tyr(NO2)-OH, Fmoc-Gly-OH, Fmoc-Ser(OtBu)-OH, Fmoc-Gln(Trt)-OH, Fmoc-Leu-OH, Fmoc-Thr(tBu)-OH, Fmoc-Val-OH, Fmoc-Ser(OtBu)-OH, and Boc-Abz-OH with the final N-terminal Boc group being left on the peptide. The peptide was then cleaved off the resin using the aforementioned procedure, and purified using a Vydac Si C18 RP-HPLC semi-preparative column (300 Å, 5 µM, 10 × 250 mm) and HPLC method A, with the desired peptide eluting at 16.2 minutes. The HPLC fractions were pooled and lyophilized to produce the peptide as a yellow powder. The peptide was analyzed using HRMS (ESI) calcd for C_50_H_77_N_15_O_18_ [M + 2H]^2+^ 587.7780, found 587.7781.

#### Cloning, Expression and Purification of SARS-CoV M^pro^ and SARS-CoV-2 M^pro^

DNA encoding the main protease M^pro^ from SARS-CoV-2 was obtained from BioBasic Inc. (Ontario, Canada) and codon optimized for expression in *Escherichia coli*. The gene was cloned into the pET SUMO (small ubiquitin-like modifier) expression vector (Invitrogen). Clones were sequenced to ensure that the SARS-CoV-2 M^pro^ protein was in frame with the His-tagged SUMO fusion protein. The resulting plasmid was transformed into *E. coli* BL21(DE3) and the *E. coli* transformant was grown in Luria Broth, Miller at 37 °C with shaking (220 rpm) to an OD_600_ of 0.6-0.7 using kanamycin (50 μg/mL) as selective pressure. Expression of the fusion protein was induced by the addition of 0.5 mM IPTG to the cell culture and the culture was grown for an additional 4-5 h at 37 °C. Cells were harvested by centrifugation (6000 *g* for 10 min at 4 °C) and suspended in lysis buffer (20 mM Tris-HCl pH 7.8, 150 mM NaCl). Cells were lysed by sonication on ice and the lysate was centrifuged (17000 *g* for 10 min at 4 °C) to remove cellular debris. The supernatant was isolated and, after adding imidazole (5 mM), mixed with Ni-NTA resin (Qiagen). The mixture was loaded on a fritted column and allowed to flow by gravity at 4 °C. The resin was washed with 10 column volumes (CV) of lysis buffer containing 20 mM imidazole. The fusion protein was eluted using 2 CV of lysis buffer containing increased concentrations of imidazole (40, 60, 80, 100, 200 and 500 mM). Eluted fractions were analyzed by SDS-PAGE and those that contained the fusion protein were pooled together, dialyzed against lysis buffer containing 1 mM DTT at 4 °C and concentrated using Amicon Ultra-15 filter (Millipore) with a MWCO of 10 kDa. The fusion protein was digested by His-tagged SUMO protease (McLab, South San Francisco, CA) at 4 °C for 1-2 h to remove the SUMO tag. The cleavage mixture was added to Ni-NTA resin and loaded on a fritted column. The flow through containing SARS-CoV-2 M^pro^ was collected and analyzed by SDS-PAGE. The SARS-CoV-2 M^pro^ protein was further purified using size exclusion chromatography (G-100, GE Healthcare, 1 ml/min flow rate, 4°C) in 20 mM Tris, 20 mM NaCl, 1 mM DTT, pH 7.8. Immunoglobulin G, 166 kDa; bovine serum albumin, 67 kDa; ovalbumin, 43 kDa; and lysozyme, 15kDa were used as calibration standards. Fractions containing the SARS-CoV-2 M^pro^ protein were pooled and concentrated using Amicon Ultra-15 filter with a MWCO of 10 kDa. In addition, a sample of the fusion protein SUMO-SARS-CoV-2 M^pro^ was also purified using size exclusion chromatography and concentrated as described above. The plasmid encoding the SARS-CoV M^pro^ with an N-terminal His-tag upstream of an FactorX cleavage site was the kind gift of Dr. Michael James, which was expressed a purified according to previous protocols^3^.

### Mass Spectrometry of SARS-CoV-2 M^pro^

The mass of the free SARS-CoV-2 M^pro^ was confirmed by HR-MALDI on a MALDI-TOF (Bruker Ultrafelxtreme, Bruker Daltronics, USA) and LC-MS on an ESI-TOF instrument (Agilent Technologies 6220, California, USA) using electrospray ionization.

### Enzyme Kinetics of SARS-CoV-2 and SARS-CoV M^pro^

A synthesized fluorescent substrate containing the cleavage site (indicated by the arrow, ↓) of SARS-CoV-2 M^pro^ (2-Abz-SVTLQ↓SG-Tyr(NO_2_)-R-NH_2_) was used for the fluorescence resonance energy transfer (FRET)-based cleavage assay^4^. The protease reactions of both SARS-CoV-2 M^pro^ and SARS-CoV M^pro^ towards fluorescent substrate was performed in activity buffer (20 mM Bis Tris, pH 7.8, 1 mM DTT) at 37 °C for 10 min. The final concentration of proteases used in the assay was fixed at 80 nM and the concentrations of the substrate were varied from 0.1 to 500 μM. Reaction was started with the enzyme and the fluorescence signal of the Abz-SVTLQ peptide cleavage product was monitored at an emission wavelength of 420 nm with excitation at 320 nm, using an Flx800 fluorescence spectrophotometer (BioTek). Before kinetic calculations, it was verified that the proportionality between the fluorescence emitted and the amount of the substrate used in the assay was linear. The minimal concentration of the enzyme and time of reaction that gave a linear dependence of amount of generated product with time was chosen. Initial velocities in corresponding relative fluorescence units per unit of time (ΔRFU/s) were converted to the amount of the cleaved substrate per unit of time (μM/s) by fitting to the calibration curve of free Aminobenzoyl-SVTLQ. All data are corrected for inner filter effects by an adopted literature protocol. In short, the fluorescence signal (RFU) at each substrate concentration was determined and defined as f(FRET). Then, 5uL free Aminobenzoyl-SVTLQ at final 5uM was added to each concentration and fluorescence was taken f(FRET+ Aminobenzoyl-SVTLQ). Simultaneously, a reference reading was taken with the same free Aminobenzoyl-SVTLQ concentration and defined as f(ref). The inner-filter correction was obtained as: corr% = (f (FRET + Aminobenzoyl-SVTLQ) – f (FRET)) / f (ref) x 100% The corrected initial velocity of the reaction was calculated as

V = V_o_ / (corr%).

V_o_ represents the initial velocity of each reaction.

Kinetic constants (v_max_ and K_m_) were derived by fitting the corrected initial velocity to the Michaelis-Menten equation, v = v_max_ × [S] / (K*_m_* + [S]) using GraphPad Prism 6.0 software. k_cat_/K*_m_* was calculated according to the equation, k_cat_/K_m_ = v_max_ / ([E] x K_m_). Triplicate experiments were performed for each data point, and the value was presented as mean ± standard deviation (SD).

### Inhibition parameters

Stock solutions of GC373 and GC376 were prepared with 10% aqueous DMSO. For the determination of the IC_50_, 80 mM of SARS-CoV-2 M^pro^ was incubated with GC373 or GC376 at various concentrations from 0 to 100 μM in 20 mM Bis-Tris, pH 7.8, 1 mM DTT at 37°C for 10 min. The protease reaction was started by addition of 100μM of the substrate. The GraphPad Prism 6.0 software (GraphPad) was used for the calculation of the IC_50_ values. Both inhibitors were tested for non-specific binding by performing a reference titration in the absence of DTT showing no influence in the obtained fluorescence readings (data not shown).

### Crystallization and Structural Determination

#### Crystallization

For crystallization, purified SARS-CoV-2 M^pro^ was dialysed against buffer containing 10 mM NaCl and 5mM Tris HCl pH 8.0 overnight at 4 °C, and concentrated with a Millipore centrifugal filter (30 kDa MW cutoff) to a concentration of 9 mg/mL. Protein was incubated with 5 molar excess of inhibitor at 4 °C for 2 h prior to crystallization. For SARS-CoV-2 M^pro^, the protein was subjected to the PACT crystallization screen (Molecular Dimensions), with hits identified in several conditions for both inhibitors. Best crystals were observed with hanging drop trays at room temperature at a ratio of 1:1 with mother liquor 0.2 M Sodium sulfate, 0.1 M Bis-Tris propane pH 6.5, 20 % w/v PEG 3350. While the SARS-CoV-2 M^pro^ with ligands crystallize with mother liquid containing 0.2 M Sodium chloride 0.1 M HEPES pH 7.0 20 % w/v PEG 6000. Prior to freezing, crystals were incubated with 15% glycerol as a cryoprotectant. Crystals were initially screened on our 007 MicroMax (Rigaku Inc) home source with final data collection at SSRL, beamline 12-2.

#### Diffraction Data Collection, Phase Determination, Model Building, and Refinement

All diffraction data sets were collected using synchrotron radiation of wavelength 0.97946 Å at beamline 12-2 of Stanford Synchrotron Radiation Lightsource (SSRL) California, USA, using a Dectris PILATUS 6M detector. Several data sets were collected from the crystals of SARS-CoV-2 M^pro^ free enzyme as well as with GC376 and GC373 treated. XDS^5^ and Scala were used for processing the datasets. The diffraction dataset of the free SARS-CoV-2 M^pro^ was processed at a resolution of 1.75 Å, in space group P2_1_ (Table S1). For the complex of SARS-CoV-2 M^pro^ with GC376 and GC373, the dataset collected, was processed at a resolution of 1.9 Å and 2.0 Å and in space group C2 (Table S1). All three structures were determined by molecular replacement with the crystal structure of the free enzyme of the SARS-CoV2 M^pro^ (PDB entry 6Y7M as search model, using the Phaser program from Phenix. Ligand Fit from Phenix was employed for the fitting of both inhibitors in the density of pre-calculated map from Phenix refinement, using the ligand code K36. Refinement of the three structures was performed with phenix.refine in Phenix software. Statistics of diffraction, data processing and model refinement are given in (Table S1).

### GC373 NMR Binding Assay

#### NMR Samples

NMR samples were first prepared by dialyzing SARS-CoV-2 M^pro^ enzyme to exchange buffers (target buffer: D_2_O, 50 mM phosphate, pD 7.5 with 20 mM DTT) by spin filtration (Amicon micro-spinfilter, 10 kDa cutoff). A 50 µL solution of 2.6 mg/mL enzyme was added to the spin filter and diluted to 300 µL, then spun at 6600 g for 18 min. This was repeated an additional 2 times, and the sample was then made up to 300 µL in volume and transferred to the NMR tube. Samples of enzyme in the presence of inhibitor were prepared by administering an additional 1.5 µL of ^13^C labelled GC373 solution (20 mM in DMSO) to the aforementioned enzyme sample. Sample data was acquired in 5 mm Varian specific D_2_O susceptibility matched microcell (i.e. BMS-00V) Shigemi NMR tubes purchased from Wilmad Lab-glass Inc. All NMR solvents were purchased from Sigma Aldrich. NMR tubes were washed between runs using 5 rinses of D_2_O, and then inverted to air dry overnight.

Small volume additions (e.g. inhibitor added to enzyme) to samples in Shigemi NMR tubes were done by adding to the top of the outer NMR tube with the inner plunger removed, and then carefully tapping the sample tube in an almost horizontal position while rotating to break the surface tension and allow sample liquid to flow up the tube. The sample volume was allowed to travel up until making contact with the additional material. The tube was then repeatedly whipped downward to move the material to the bottom. This was repeated several times so that additions were rinsed down into the microcell ensuring proper mixing.

#### NMR Spectroscopy

NMR experiments were collected at 16.45 T (i.e. 700 MHz) using a 4-channel ‘VNMRS’ (Varian/Agilent) NMR spectrometer (VNMRJ 4.2 patch110 software) with an Agilent 7620 automatic sample handling system. A 5mm triple resonance cryogenically cooled (20 K) ^1^H direct-detection (i.e. with ^13^C^15^N on the outer coil) probe was utilized for all experiments. The probe had cooled preamplifiers on the 1H and 13C detection channels. The sample temperature was calibrated to 27°C using methanol^6^.

All spectra were run “locked” on the ^2^H resonance signal and chemical shifts were referenced using the residual proton ^1^HOD signal position^7^ (i.e. 4.7 ppm) prior to saturation. One dimensional ^1^H data were acquired using presaturation^8, 9^ for residual ^1^HOD solvent suppression, followed by a 90° excitation pulse and data acquisition. The saturation carrier position amplitude were manually optimized using a position sweep array and based on the calibrated high-power pulse, respectively. Saturation was applied directly on the water resonance (i.e. depending on water suppression efficacy and the ability to avoid analog to digital and/or receiver overloads with sufficient gain for data acquisition). The saturation was applied with a gammaB_1_ induced field strength of ∼80 Hz^10^, and a duration of 2 seconds. The 90° pulse width was determined after tuning and matching each sample, using one-pulse nutation optimization^11^. Other specific 1D acquisition parameter settings were: sweep width of 14044 Hz, acquisition time 2.5 seconds, with 70224 total (i.e. real plus imaginary) data points.

A gradient-selected phase-sensitive two dimensional ^1^H,^13^C-heteronuclear single quantum correlation (HSQC)^12–14^ with adiabatic inversion, refocusing and decoupling ^13^C pulses (i.e. gHSQCAD Varian/Agilent “ChemPack”, Krish Krishnamurthy) was used for all experiments. The HSQC spectra were acquired with ^1^H and ^13^C sweep widths (and recorded points) of 8389 Hz (∼12 ppm) and 38722 Hz (220 ppm), respectively (1678 total directly detected and 32 complex indirectly detected points, also respectively). A gradient stabilization delay of 500 µs was used and 146 Hz J_HC_ applied for INEPT transfers. Adiabatic inversion/recovery ^13^C pulses were applied at a 13.1 kHz induced field with a 600 us duration covering ∼400 ppm. Adiabatic decoupling was applied at ∼2.6 kHz induced field during the entire 100ms acquisition period. Carbon chemical shift referencing and carrier position (100 ppm) was based on indirect IUPAC ^1^H referencing^15^. The carbon carrier position was moved (e.g. 80, 90, or 100 ppm) to eliminate the possibility of peaks of interest near the carrier position being artifactual (i.e. quadrature “glitch”).

For processing of NMR data, all dimensions were zero-filled to twice the number of acquired points. A line-broadening apodization function of 0.25 Hz was applied for ^1^H-1D spectra while a π/2 squared sine-bell weighting function was utilized for the 2D-HSQC (both dimensions). No linear prediction was applied nor was non-linear/non-uniform data acquisition utilized to avoid any possible artifacts in the resulting data. The final spectra were manually phased and baseline corrected.

### Determination of EC_50_ by Plaque Assay

SARS-CoV-2/CANADA/VIDO 01/2020 was a kind gift from Darryl Falzarano (University of Saskatchewan). Vero (Female green monkey kidney) E6 cells were infected with an MOI of .0001 pfu/cell in infection medium consisting of DMEM supplemented with 1x non-essential amino acids (Gibco), 10 mM HEPES, 2% fetal bovine serum, 50 IU/mL penicillin, 50 IU/mL streptomycin different doses of antiviral drugs. After 1h, the infecting medium was removed and monolayers were overlaid with MEM supplemented with 10mM HEPES and 1.2% Avicel RC-591 (DuPont). After 48 h, cells were fixed in 10% formaldehyde, and stained using 0.5% (w/v) crystal violet. Plaques were counted and data was plotted as % inhibition vs the log_10_[drug] using Prism (GraphPad). EC50’s were determined using a non-linear regression analysis. Experiments were done in triplicate. Error bars indicate standard deviation.

### Quantification of SARS-CoV-2 Viral RNA in Cell Culture Supernatants by qRT-PCR

Cell supernatants (140 µL) were collected at various points after infection, and RNA was isolated using the QIAmp Viral RNA Mini kit as per manufacturer’s instructions (Qiagen). Reverse transcription was carried out on 2 µL using Superscript IV Vilo master mix (Invitrogen). Quantitative PCR was carried out using 2 µL of cDNA in TaqMan Fast Master mix using primers and probe for the N gene (N2 primers) designed by the United States center for disease control and prevention (IDT cat#10006606). A standard curve was generated using dilutions of positive control standards from CDC (IDT cat # 10006625).

### Measuring Cytotoxicity in A549 and Vero E6 cells

Cell viability was measured using the CellTiter-Glo luminescent cell viability assay (Promega). Either A549 (male human lung epithelial) cells or VeroE6 cells were seeded at 5×10^3^ cells/well in 96-well plates and incubated overnight before treatment. Compounds GC373 and GC376 were solubilized in DMSO and added to cells in an eight-point four-fold serial dilution (200 µM to 0.0122 µM). Cells were incubated in the presence of compounds for 24 hours before addition of the luminescence substrate and measurement of ATP activity according to manufacturer’s instructions. The percentage of viable cells was calculated relative to cells treated with solvent alone (0.5% DMSO). Results were plotted as the mean of three independent experiments ±SD, where each experiment consisted of quadruplicate wells per concentration of compound.

## NMR Spectra of Synthesized Compounds

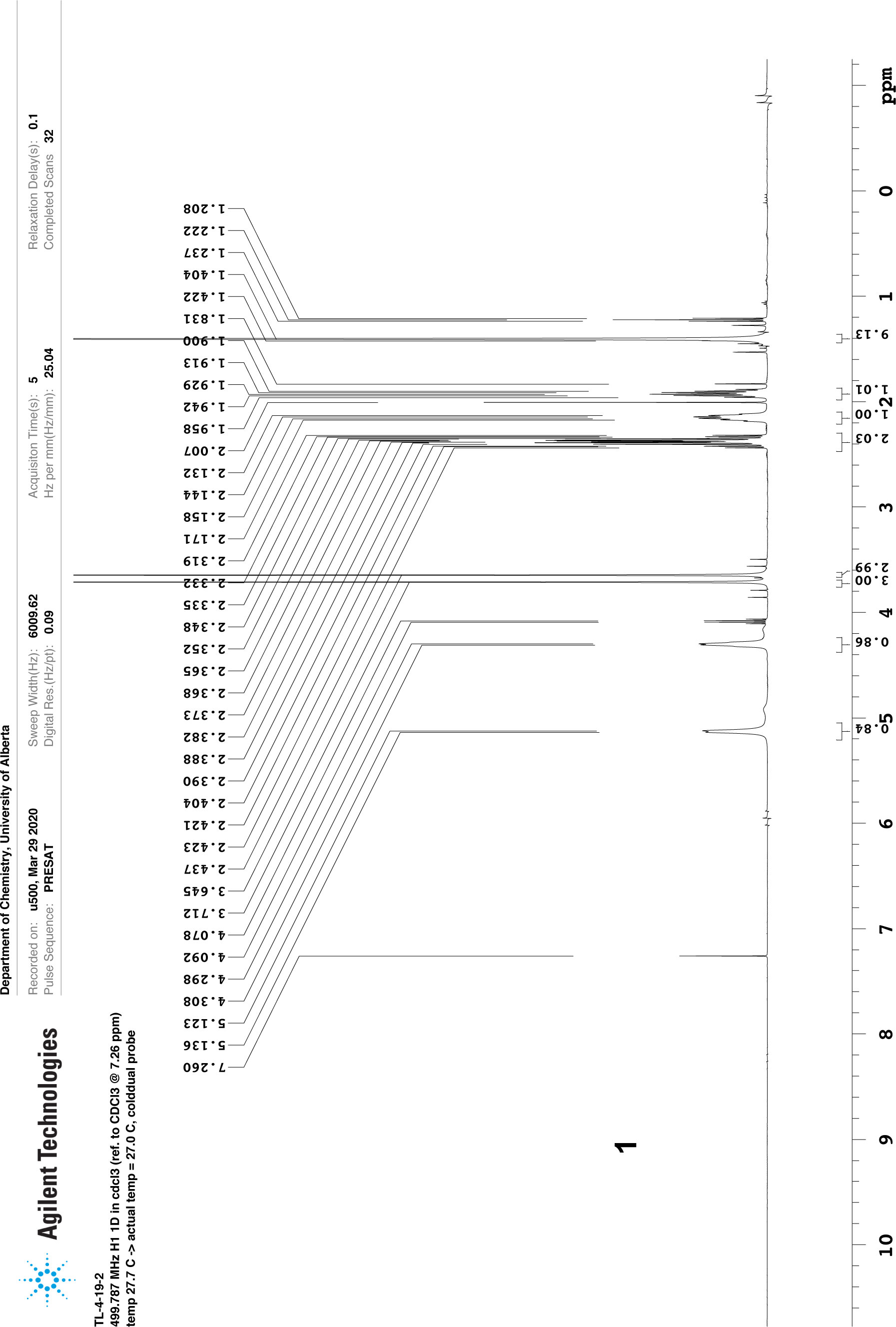
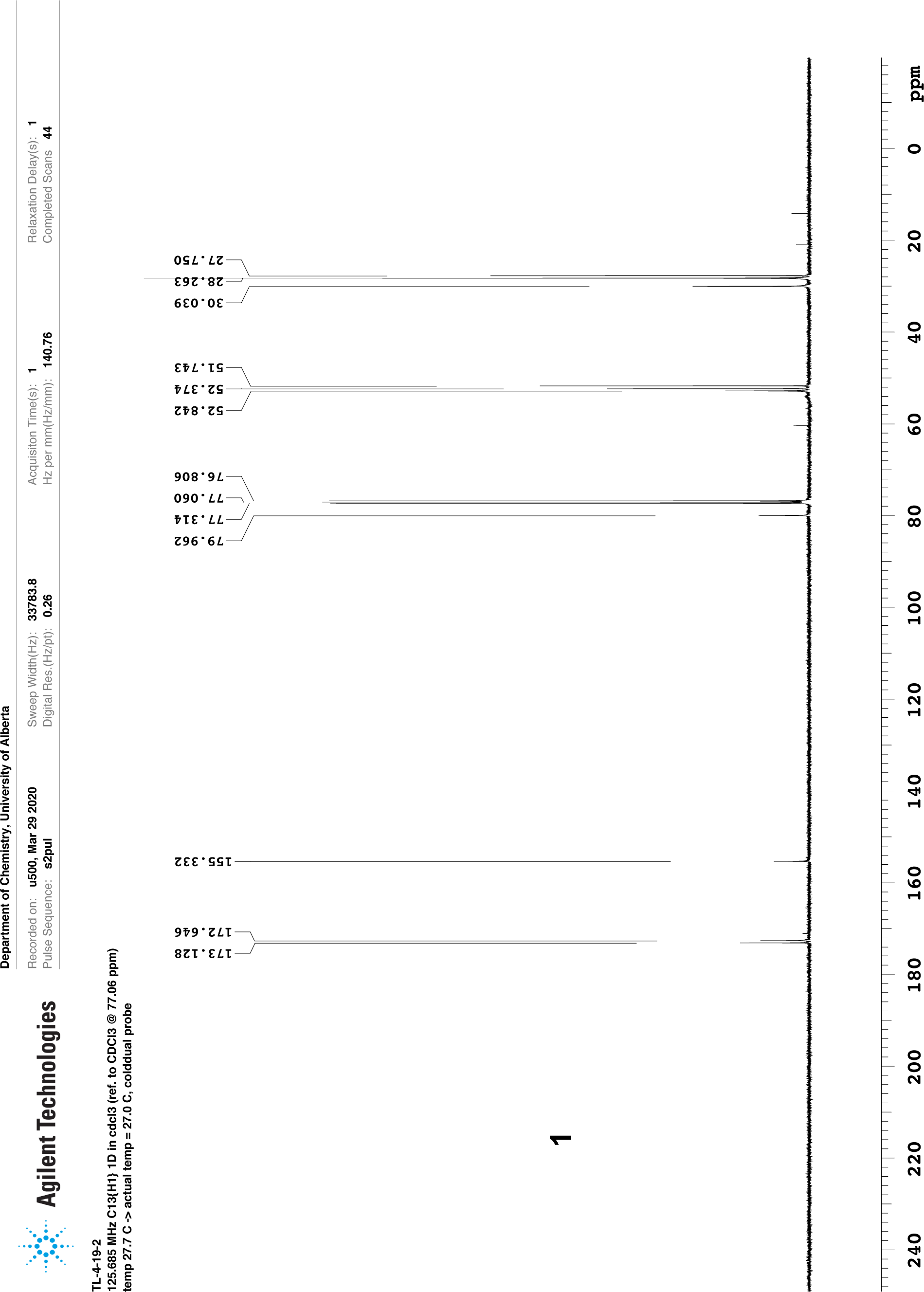
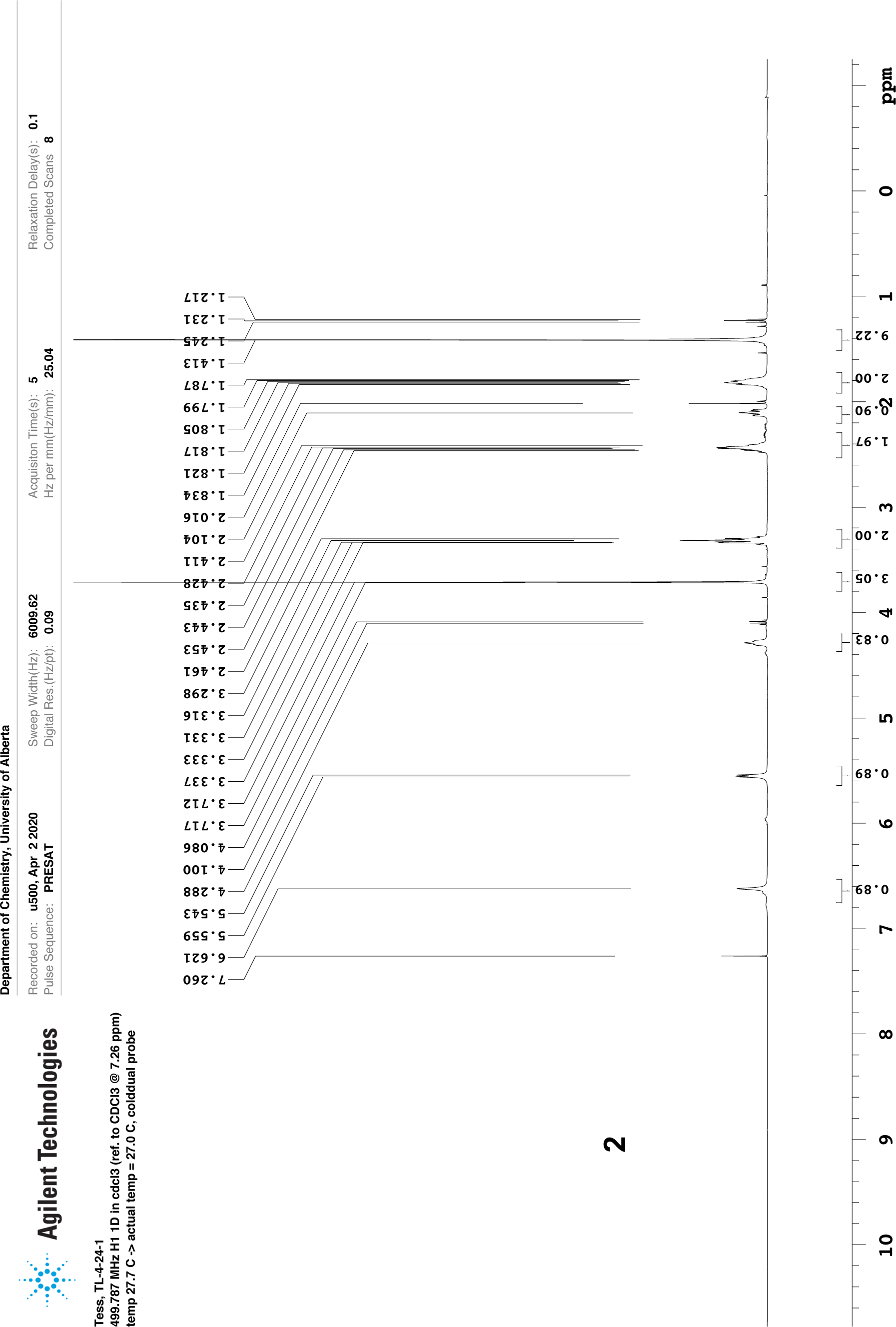
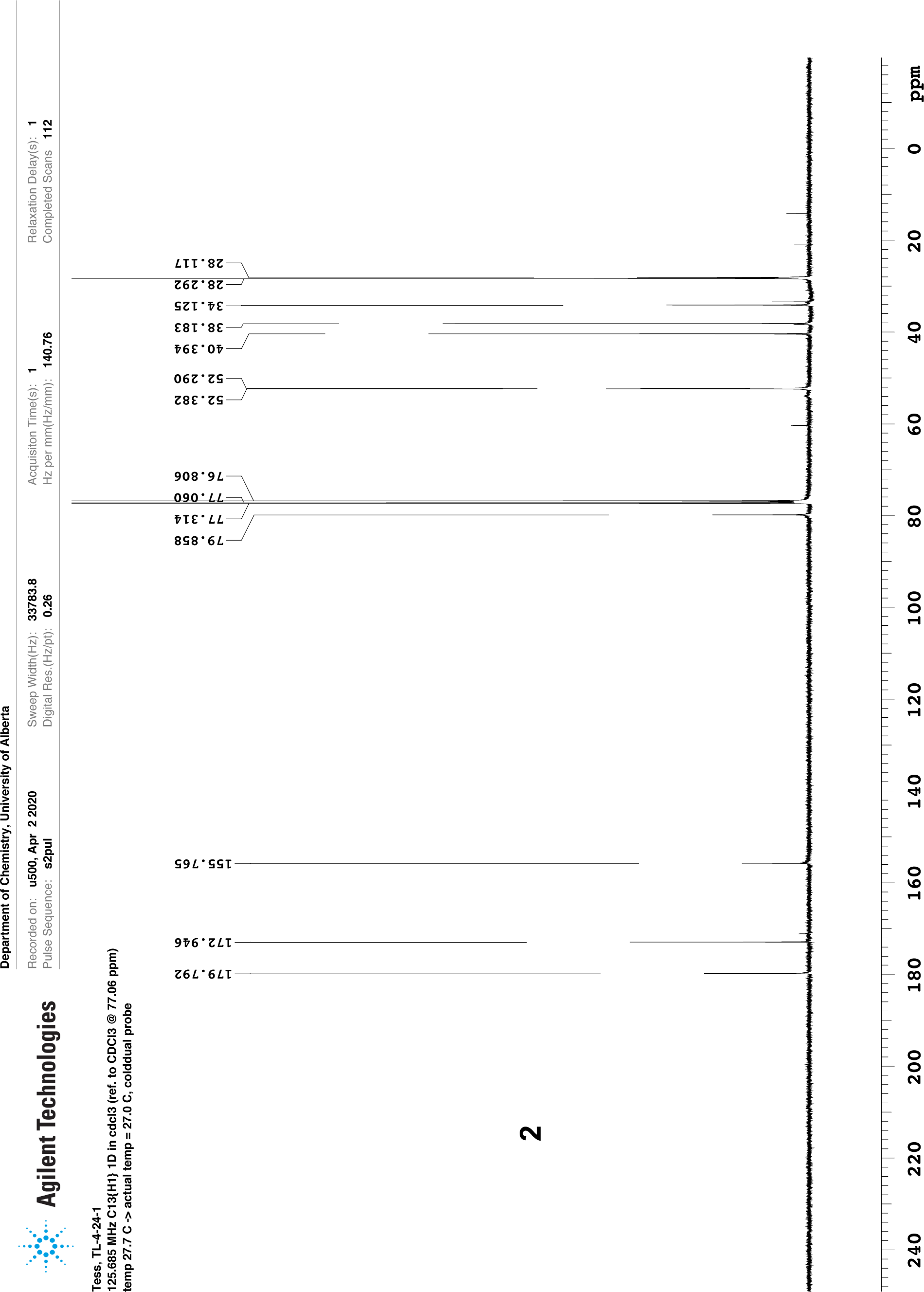
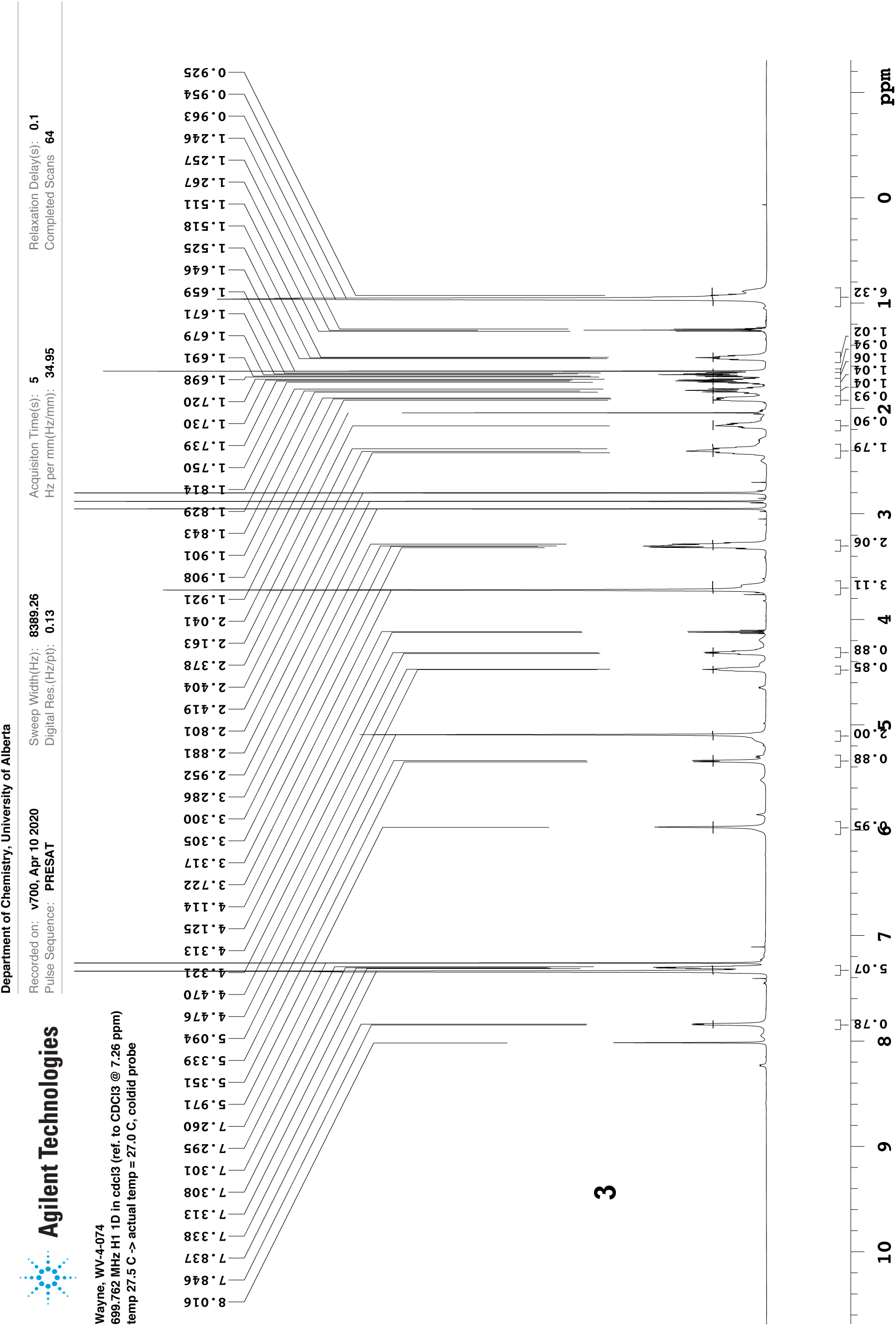
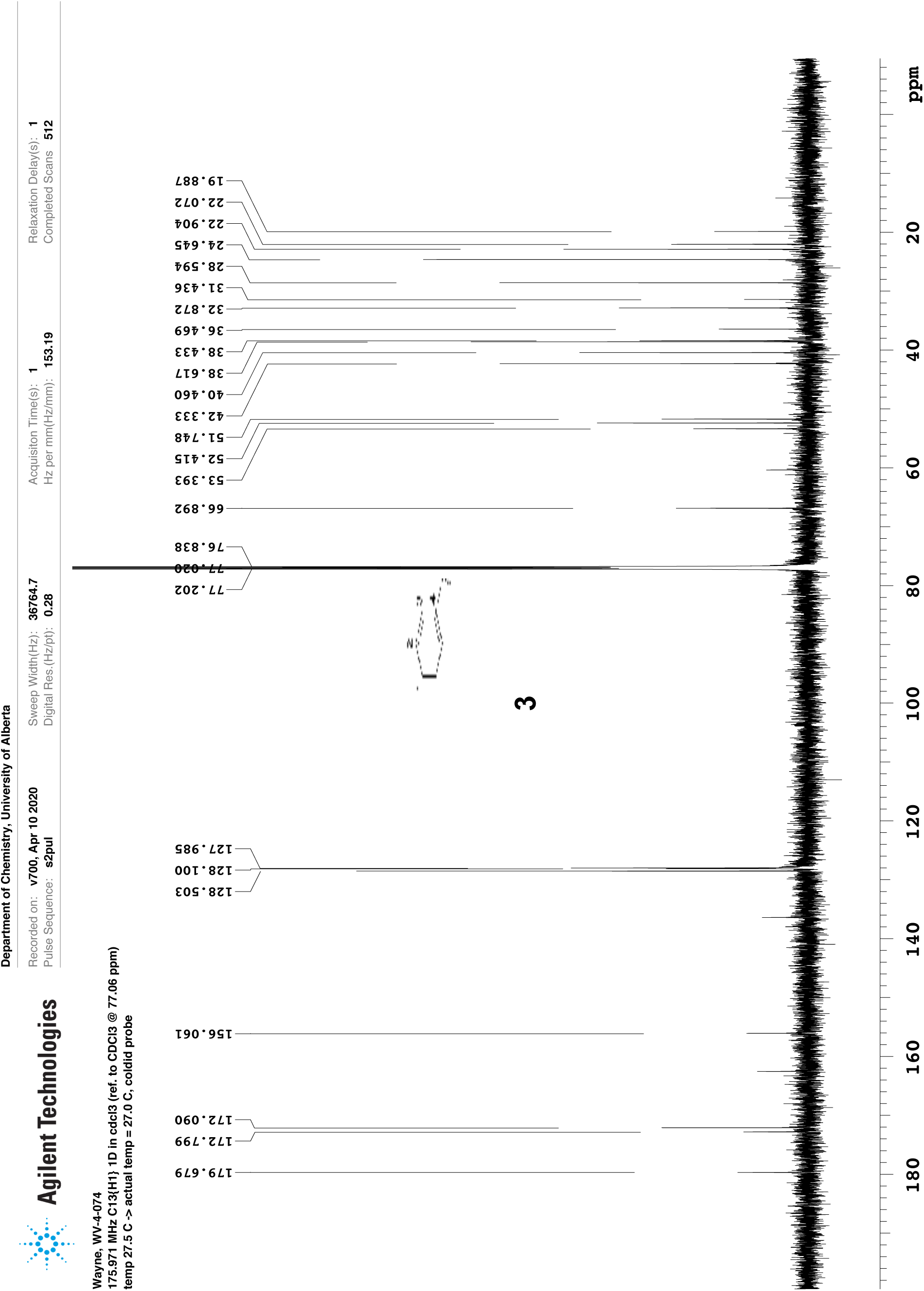
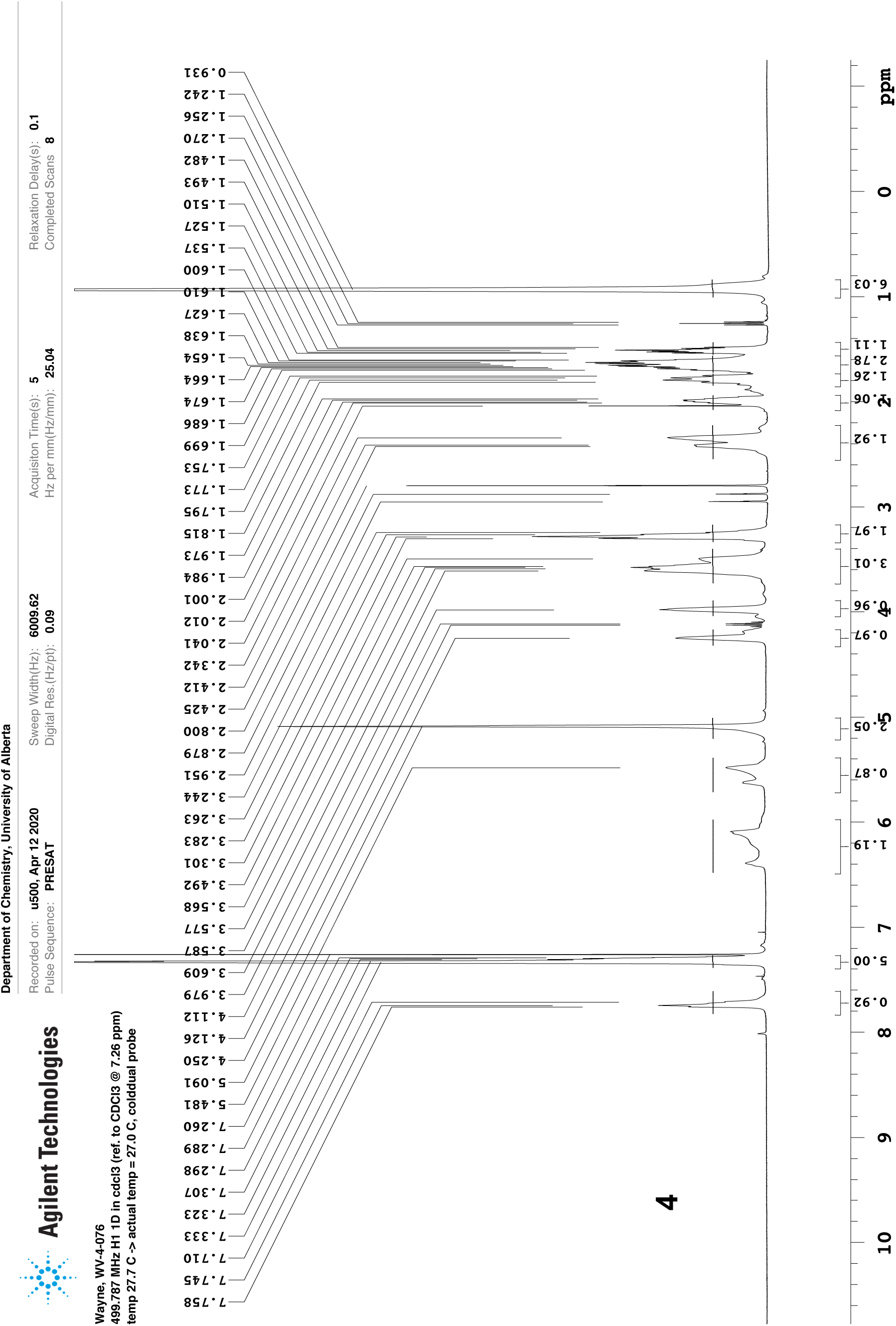
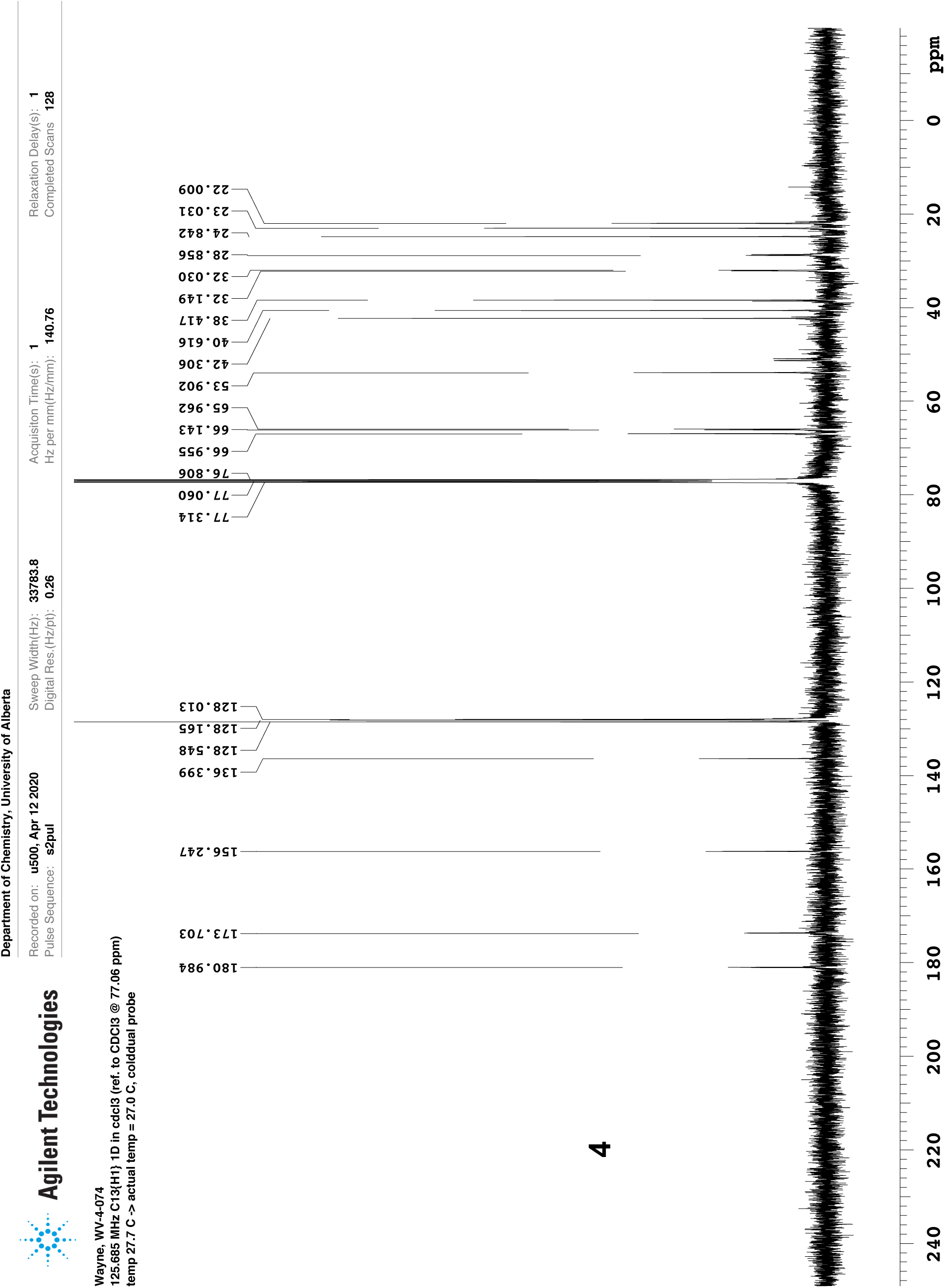
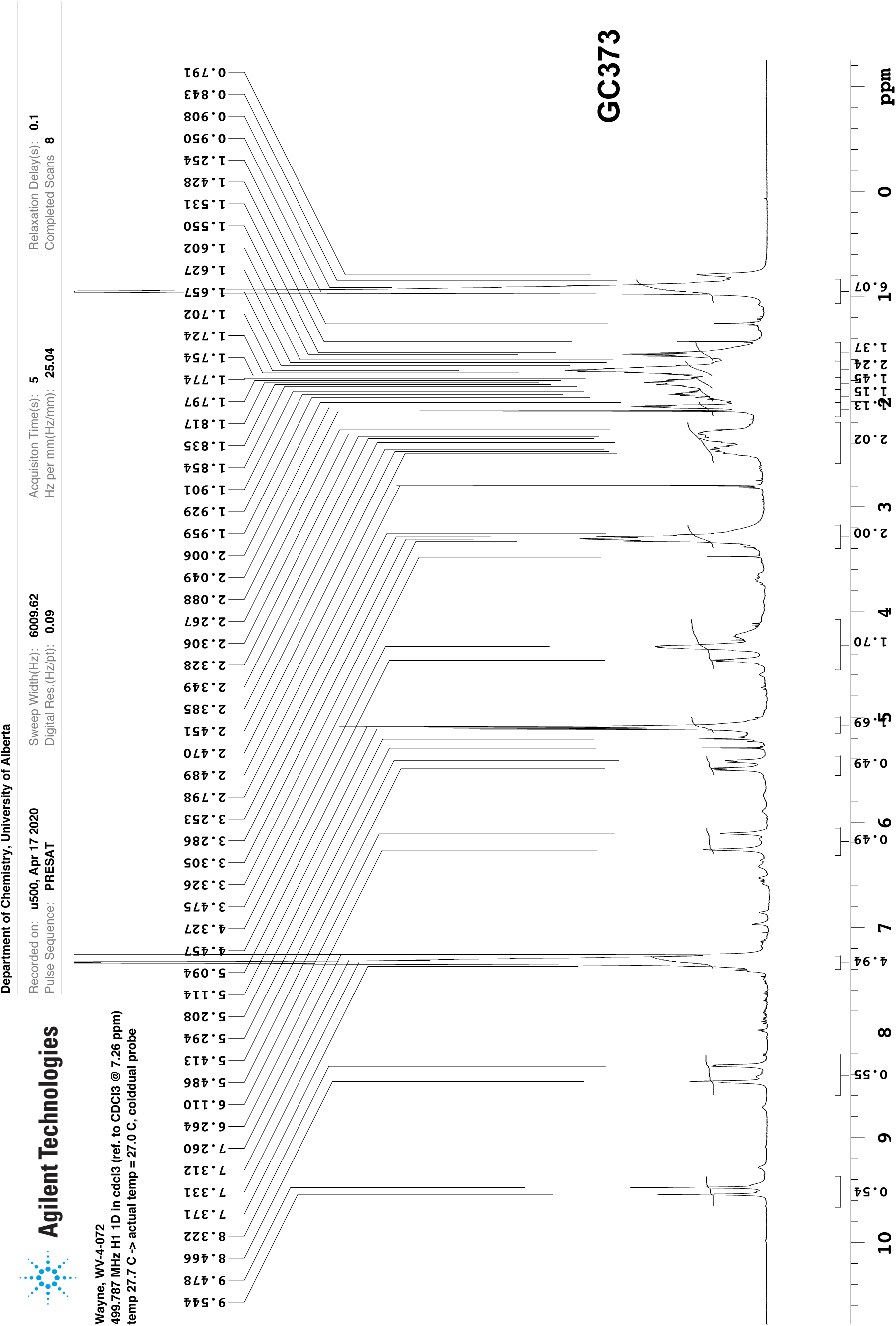
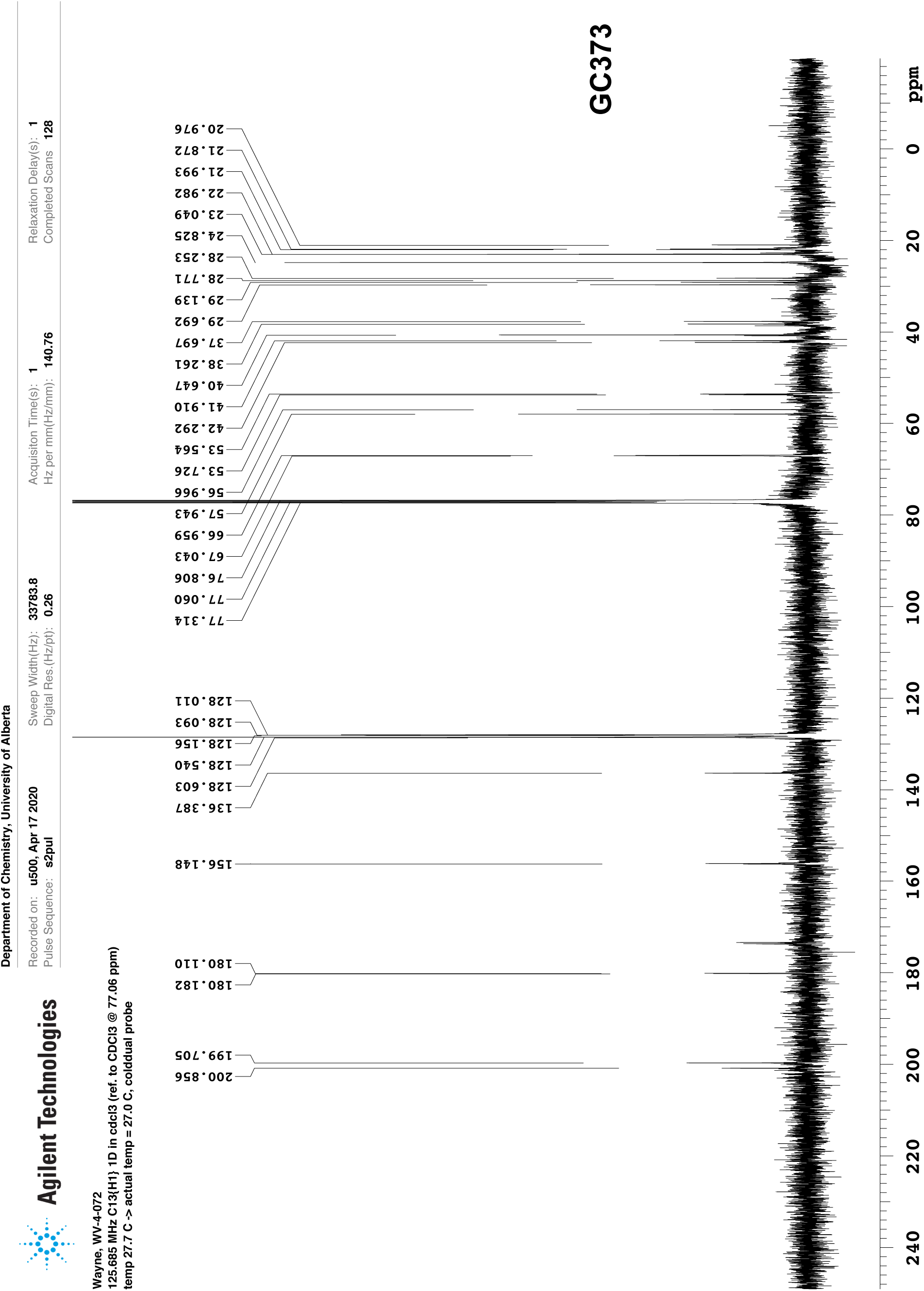
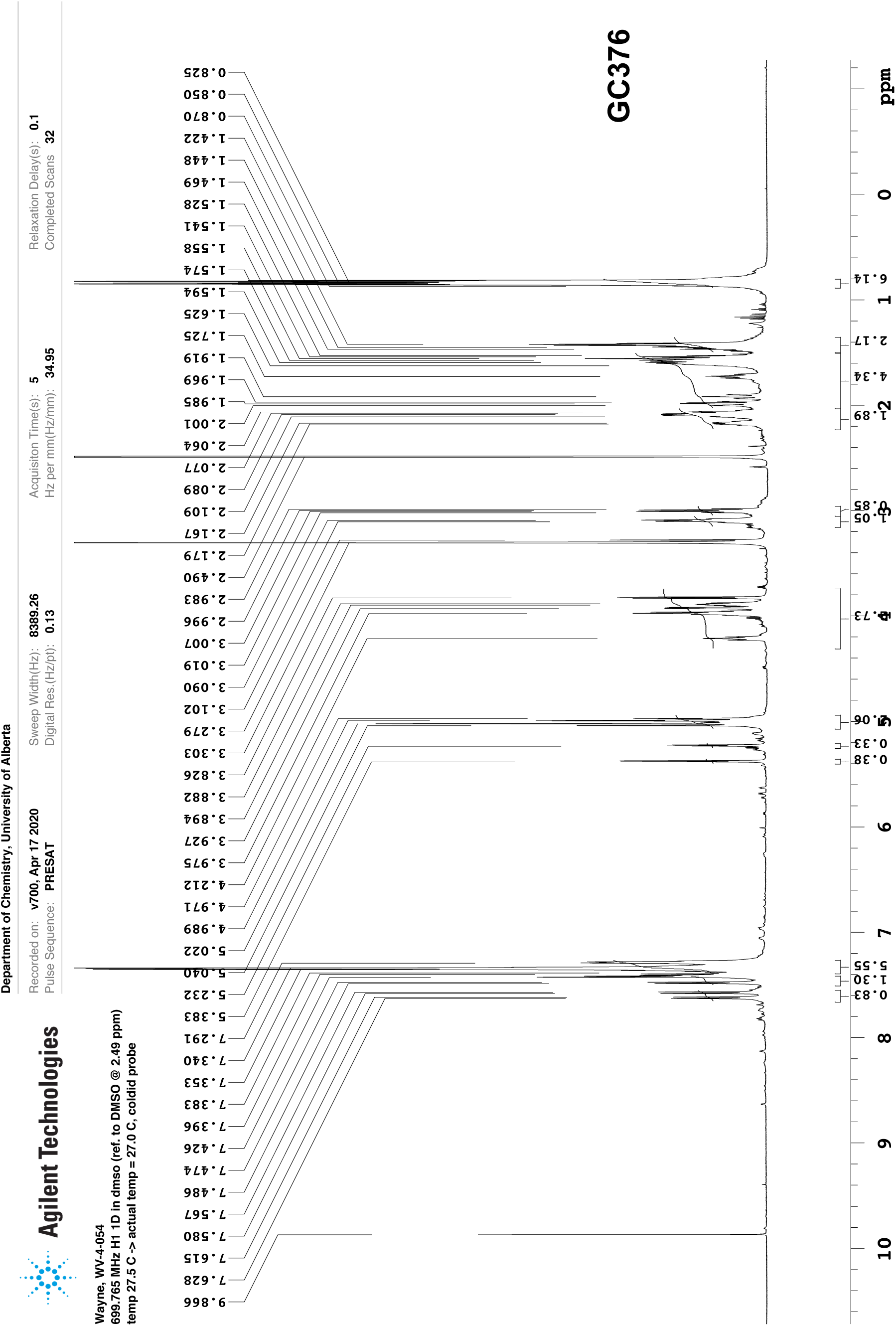
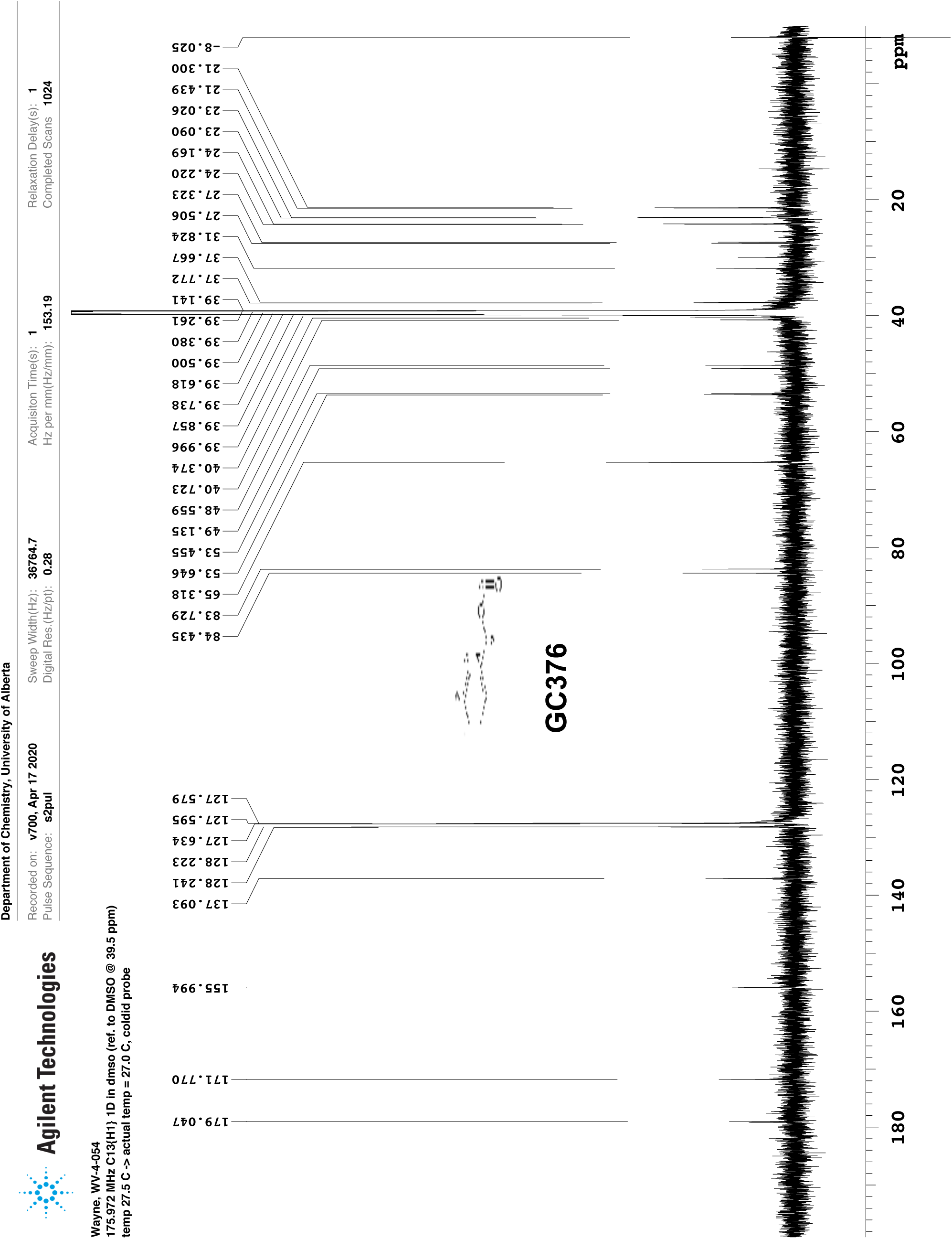

